# Comparative transcriptomics uncovers plant and fungal genetic determinants of mycorrhizal compatibility

**DOI:** 10.1101/2025.04.11.648352

**Authors:** José Eduardo Marqués-Gálvez, Maira de Freitas Pereira, Uwe Nehls, Joske Ruytinx, Kerrie Barry, Martina Peter, Francis Martin, Igor V Grigoriev, Claire Veneault-Fourrey, Annegret Kohler

**Author notes:** Both authors equally contributed as first author (José Eduardo Marqués-Gálvez, Maira de Freitas Pereira).

## Abstract

Ectomycorrhizal symbiosis supports tree growth and is crucial for nutrient cycling and temperate and boreal ecosystems functioning. The establishment of functional ECM first requires the association of compatible partners. However, host and fungal genetic determinants governing mycorrhizal compatibility are unknown. To identify such factors in poplar and its fungal associates, we mined existing and *de-novo* tree and fungal transcriptional datasets. We identified core plant regulons enabling ECM symbiosis at early and mature stages of the interaction. These regulons can be divided into general fungal-sensing and ECM-specific components. We highlight the importance of fungal modulation of plant JA-related defences and the regulation of secretory pathways for ECM compatibility, including upregulation of key fungal small secreted proteins, the downregulation of plant secreted peroxidases and the downregulation of plant cell-wall remodelling proteins concomitantly with the upregulation of fungal glycosyl hydrolases acting on pectin. Not only gene-regulation, but also its temporal scale and dynamics seems to play a crucial role for mycorrhizal compatibility. The expression profile of the host Common Symbiosis Pathway was also studied, revealing constitutive levels of expression of a part of the pathway and moderate upregulation in compatible ECM interactions. Overall, these results underscore the importance of novel biological functions during the establishment of ECM symbiosis, help us gain insights into the molecular events determining mycorrhiza compatibility and serve as a data-rich transcriptomic resource to open new research questions in the field.

**Significance statement:** Ectomycorrhizal symbiosis is essential for forest ecosystems, but the genetic determinants of host and fungal compatibility remain underexplored. Employing comparative transcriptomics, we identified plant general and ECM-specific gene regulons, outlining the expression profile of Common Symbiosis Pathway genes in multiple ECM interactions and highlighting fungal and plant genes associated to interkingdom crosstalk and cell wall remodelling. These findings help us gain insights into the molecular events determining mycorrhiza compatibility.

## Introduction

Plant roots interact with a multitude of soil microorganisms, that can range from detrimental to beneficial. Plant-microbe mutualistic interactions include mycorrhizal symbiosis, which has been commonly divided in four different main groups: Arbuscular Mycorrhiza (AM), Ectomycorrhiza (ECM), Orchid Mycorrhiza (OM) and Ericoid Mycorrhiza (ERM). Among them, ECM symbiosis plays a predominant role in temperate and boreal forests, where they participate in nutrient cycling and carbon sequestration. It has been estimated that there are around 20,000 different species of ECM fungi, including Ascomycota, Basidiomycota and Mucoromycota. These fungi interact with around 6,000 plant species, mostly trees and shrubs belonging to Pinaceae and most Angiosperms that commonly dominate temperate forest ecosystems (Tedersoo et al., 2012; van der Heiden et al., 2015; Brundrett and Tedersoo, 2018; Martin et al., 2024). Although ECM symbioses have independently appeared between diverse fungal and plant lineages, they share a common anatomical pattern of root colonization, representing an outstanding example of convergent evolution in multispecies interactions (Bittleston et al., 2016). These common morphological traits consist in hyphae aggregation that surrounds the lateral root tips of trees (i.e., mantle) and *in planta* fungal colonization of the apoplastic spaces between the root cortical cells (i.e., Hartig net), which serves as the nutrient exchange surface between the tree and the fungus (Peterson and Massicote, 2004). It thus represents a good indicator of host-fungus compatibility and is correlated with host growth responses (Barker et al., 1998; Smith and Read, 2010).

The compatibility between host tree and ECM fungus exhibits a wide range of spectrum. Some trees form associations with many different fungi, but others interact with a moderate number of species (Molina and Trappe, 1982; Liao et al., 2016). On the other side, some fungi present narrow host specificity (host specialists), while others have broad host range (host generalists) (Molina and Trappe 1982). While the molecular events resulting in successful ECM symbiosis are being increasingly studied, not much is known about the host and fungal genetic determinants of mycorrhizal compatibility. Recent advances in plant genomics and transcriptomics have boosted the discovery and characterization of gene-regulatory networks that contribute to our general understanding of ECM compatibility. The recent availability of ECM-establishing tree reference genomes (Tuskan et al., 2006; Nystedt et al., 2013; Myburg et al.,2014; Salojärvi et al., 2017; Plomion et al., 2018) led to the identification of core transcriptional programs which support the existence of common ECM symbiotic pathways in *Quercus robur* (Bouffaud et al., 2020), or provide new perspectives for the plant Common Symbiosis Pathway (CSP) (Marqués-Gálvez et al., 2022 and references therein), an ancestral pathway used by both AM fungi and nitrogen-fixing rhizobia bacteria (Olroyd, 2013). For instance, *CASTOR/POLLUX* and a calcium/calmodulin-dependent protein kinase (*CCaMK*) genes, which are important members of the CSP in *Populus trichocarpa*, seem to be involved in the interaction with the ECM fungus *Laccaria bicolor* (Cope et al., 2019). In parallel to their hosts, advances in fungal comparative genomics have also shed light over convergent fungal genetic traits related to ECM symbiosis (Martin et al., 2008, 2010; Kohler et al., 2015; Peter et al., 2016; Murat et al., 2018; Miyauchi et al., 2020; Marqués-Gálvez et al., 2021; Lofgren et al., 2021; Wu et al., 2022; Lebreton et al., 2022; Looney et al., 2022; Maillard et al., 2023; Plett et al., 2023). These studies highlight the importance of fungal secretomes, supporting that the loss of Carbohydrate-Active enzymes (CAZymes) and the expansion of small secreted effector-like orphan encoding genes upregulated during establishment of the symbiosis, named as Mycorrhizal-induced Small Secreted Proteins (MiSSPs), are a common denominator of ECM lifestyle for fungi, regardless their lineage (Kohler et al., 2015; Pellegrin et al., 2015; Miyauchi et al., 2020). A study of gene expression patterns in several compatible and incompatible *Pinus* spp. x *Suillus* spp. interactions revealed the expression of plant and fungus unique gene sets during incompatible vs. compatible pairings, highlighting fungal G-protein signalling and secretory pathways, and plant leucine-rich repeat and pathogen-resistance proteins as main actors of ECM compatibility (Liao et al., 2016). Enhanced host specificity of *Suillus* spp. also related to a significant enrichment in secondary metabolites (Lofgren et al., 2021). However, these studies are limited to specific host-partner interactions and more comprehensive studies involving different fungal and host species are needed to identify the transcriptional determinants dictating mycorrhizal compatibility, from both partners.

Here, we performed a transcriptomic analysis of seven independent interactions between poplar roots and ECM fungi, displaying compatible and non-compatible interactions. We first hypothesized that poplar roots present a core gene regulon during pre-symbiosis step and the establishment of mature ECM. We thus data-mined pre-existing and novel poplar transcriptomes at different stages of compatible ECM establishment involving different fungal clades, such as Basidiomycota (*Laccaria bicolor*, *Amanita muscaria* and *Pisolithus microcarpus*) and Ascomycota (*Cennococum geophilum*). Secondly, we hypothesized that, to be ECM-specific, the poplar genes forming this core gene regulon are not regulated in non-compatible ECM interactions. We then included in our analysis the novel transcriptomic data of poplar roots in response to two ECM Sclerodermataceae fungi unable to colonize poplar roots in our experimental conditions (*Pisolithus tinctorius* and *Scleroderma citrinum*). Finally, we hypothesized that mycorrhiza compatibility is also driven by differential gene expression of fungal partners. Thus, we analysed the fungal transcriptomic response of the Sclerodermataceae fungi, which included an ECM compatible (*P. microcarpus*) and non-compatible (*P. tinctorius* and *S. citrinum*) partners.

## Results

### Poplar establishes ECM compatible interaction with *Pisolithus microcarpus*, but not with *Pisolithus tinctorius* or *Scleroderma citrinum*

Although most of the datasets included here have been generated in previous experiments, *Populus tremula x alba* interaction with Sclerodermataceae fungi was performed *de novo* for this work (**Table S1**). *Pisolithus microcarpus* stimulated poplar lateral root formation and established a compatible ECM *in vitro*, with 14.02 ± 2.03 % of root tips being colonized (**Fig. 1**, **Table 1**). We also tested *P. tremula x alba* interaction with *P. tinctorius* and *S. citrinum*, but no visible ECM root tips were observed under stereomicroscope (**Fig. 1**, **Table 1**). Furthermore, while transversal sections of lateral root tips revealed the presence of fungal mantle and Hartig net (HN) for *P. microcarpus*, only very few mantle-like structures and no significant Hartig net were detected for *P. tinctorius* and no fungal interaction at all for *S. citrinum* (**Fig. 1 G-I**). Based on these observations, we concluded that *P. tinctorius* and *S. citrinum* were non-compatible ECM partners for *P. tremula x alba* in our experimental conditions.

**Figure 1.**
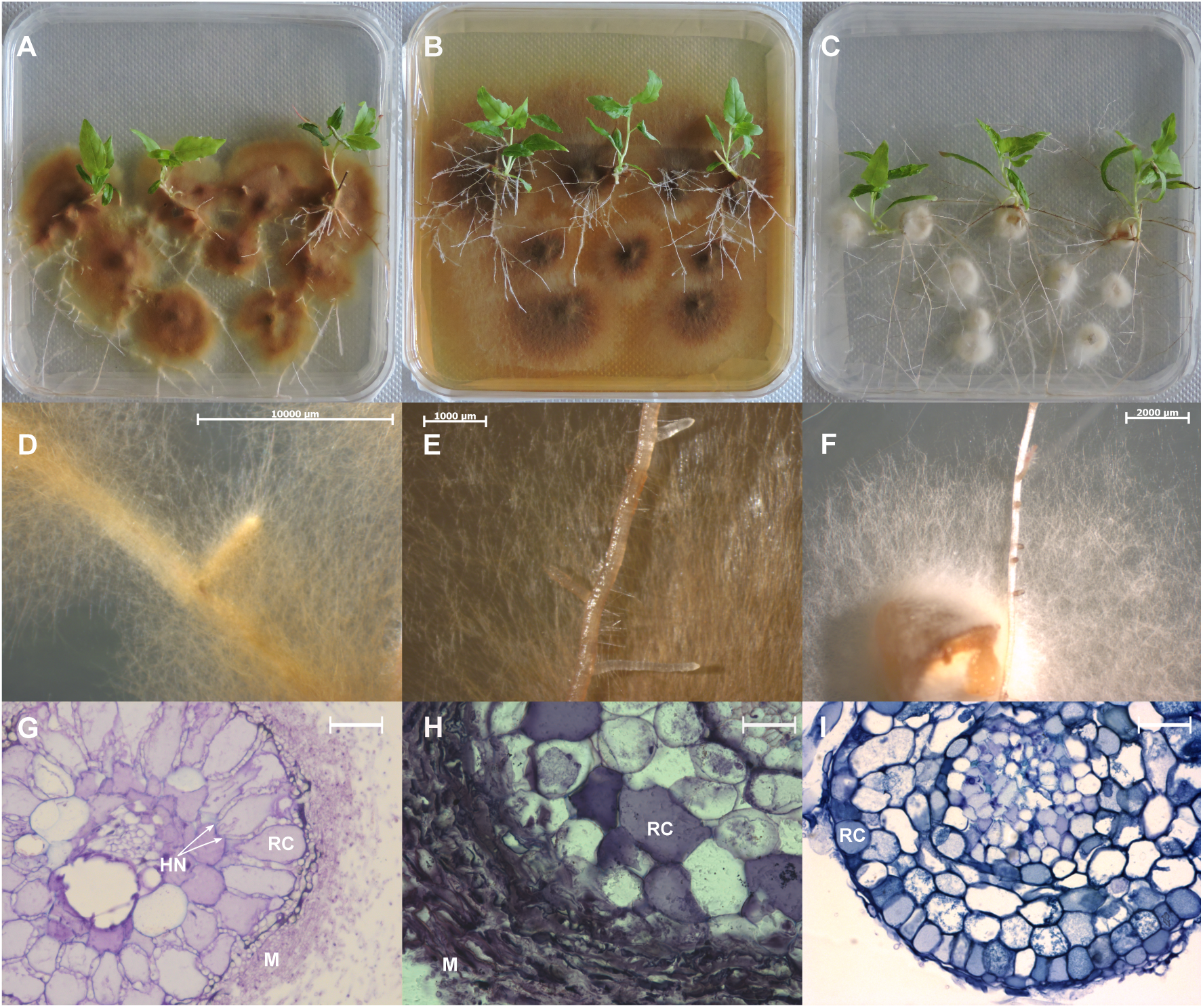
Interactions between *Populus tremula x alba* and three Sclerodermataceae fungi. The *in vitro* system in Petri dishes shows the contact between roots of *P. tremula x alba* (Pta) and (A) *P. microcarpus*; (B) *P. tinctorius*; (C) *Scleroderma citrinum*. ECM morphology with (D) monopodial branching. (E, F) Lateral root development in the *in vitro* system with no signals of ECM formation. (G) Cross-section of *P. microcarpus* x Pta ectomycorrhizas, where M = Mantle, HN = Hartig net and RC = Root cells are visible. (H) Cross-section of the interaction between *P. tinctorius* and Pta. (I) Cross-section of a lateral root produced in the presence of *S. citrinum* with no ectomycorrhizal development. The bars indicate 10000 µm (D) 1000 µm (E); 2000 µm (F) and (G-I) 50µm.

**Table 1.** Lateral root formation and ECM establishment between *P. tremula x alba* (Pta) and three Sclerodermataceae fungi. The average ± standard error is presented for each parameter. Within each time point (0-, 7-, 14- and 28-days post inoculation, dpi) and parameter (later root number and mycorrhization rate), data were submitted to Kruskal-Wallis with Fisher’s least significant difference post-hoc test. Different letters represent significant differences (p < 0.01). n = 12.

|  | 0 dpi |  | 7 dpi |  | 14 dpi |  | 28 dpi |  |
| --- | --- | --- | --- | --- | --- | --- | --- | --- |
|  | Lateral roots | Myc rate (%) | Lateral roots | Myc rate (%) | Lateral roots | Myc rate (%) | Lateral roots | Myc rate (%) |
| *Pta x P. microcarpus* | 5.50 ± 1.03 | 0 ± 0 | 17.67 ± 2.95 | 2.30 ± 1.55 | 50.83 ± 7.70 (a) | 11.44 ± 2.17 (a) | 56.42 ± 6.07 (a) | 14.02 ± 2.03 (a) |
| *Pta x P. tinctorius* | 9.03 ± 1.28 | 0 ± 0 | 20.17 ± 3.86 | 0 ± 0 | 24.17 ± 4.45 (b) | 0 ± 0 (b) | 48.67 ± 5.53 (ab) | 0 ± 0 (b) |
| *Pta x S. citrinum* | - | - | 16.5 ± 2.59 | 0 ± 0 | - | - | 39.75 ± 6.98 (b) | 0 ± 0 (b) |

### Poplar roots express a core gene regulon in response to ECM symbiosis at different developmental stages

To identify the poplar core gene regulon in response to ECM symbiosis, we performed differential gene expression analysis at two developmental stages of ECM formation (early and mature stages) from compatible interactions between *Populus* spp. and *L. bicolor*, *C. geophilum*, *A. muscaria* and *P. microcarpus*. RNAseq data coming from non-compatible *P. tinctorius* and *S. citrinum* interactions was not used for the first part of our analyses due to the lack of ECM compatibility (**Table S1**). We identified a range of 1,695 to 6,422 differentially expressed genes (DEGs, Log2 fold change > |1| and False Discovery Rate < 0.01; **Tables S2 and S3**), for single ECM compatible interactions at early or mature stages, compared to their respective non-inoculated controls (**Fig. 2A, Fig. S1**).. To identify the main drivers of the variance either technical (batch) or biological (fungal co-culture), we performed a variance partition analysis (VPA) at both early and mature stages. Our results indicated that technical variations were the main responsible of gene expression at both stages (**Fig. 2B**). Nonetheless, we found a small subset of 2,036 genes at early stage and 760 at mature stage with its variance mainly explained (≥ 50%) by the interaction with a fungus or the establishment of ECM (**Fig. 2B**). Finally, we decided to overlap the results from both differential expression and VPA analyses to only keep high-confidence genes with a significant difference in expression between interaction and control conditions and showing little variation within the interaction condition. We identified 125 and 138 genes as members of the poplar ECM core gene regulon at early and mature stages, respectively (**Fig. 2C**, **Table S4**), namely “early ECM core gene regulon” and “mature ECM core gene regulon”. Both early and mature ECM core gene regulons were mainly composed of predicted cytoplasmic globular proteins, and roughly a third (28.8% and 34.1%, respectively) of predicted secreted proteins (**Fig. 2C**).

**Figure 2.**
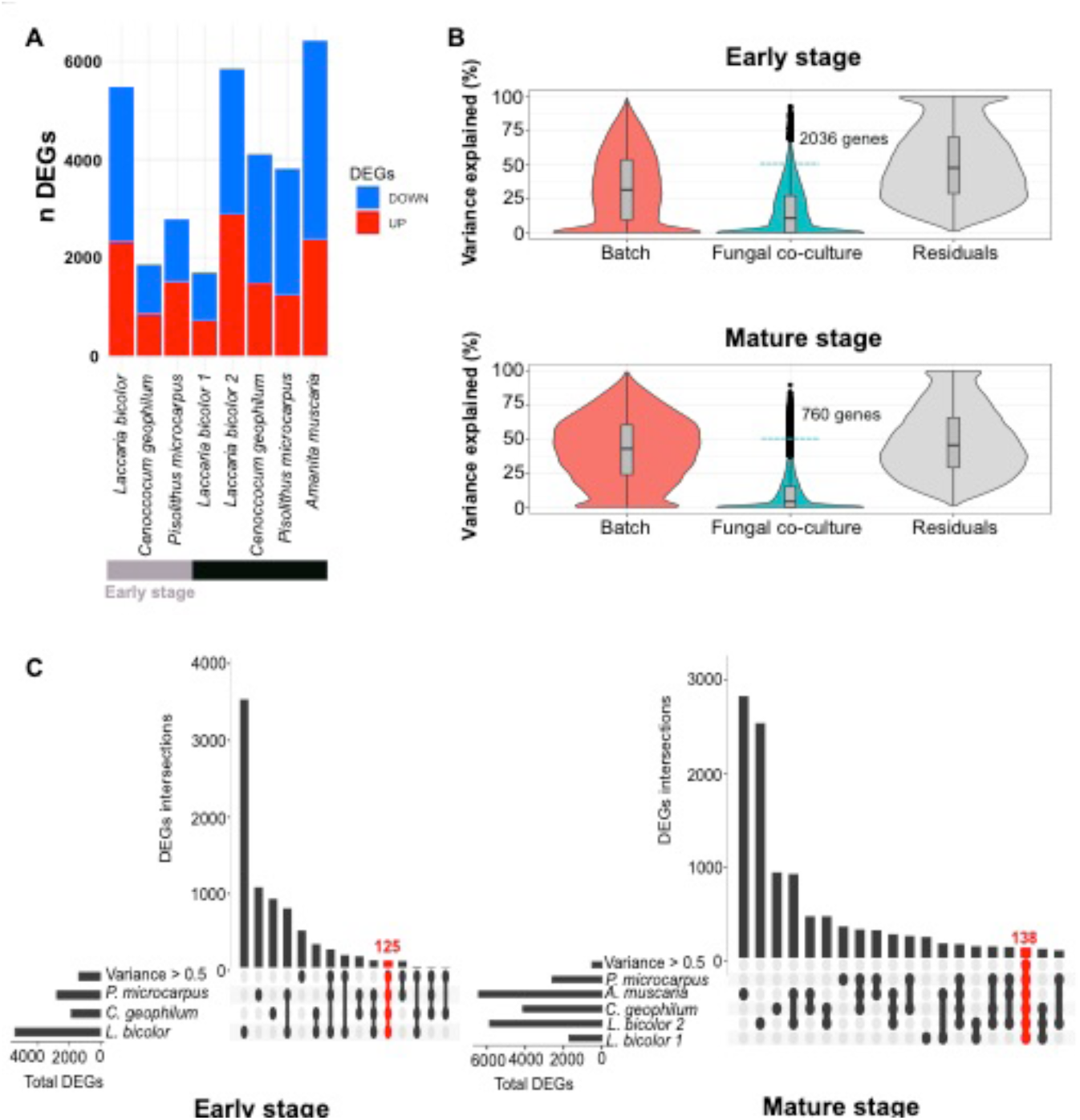
Transcriptomic analysis of poplar roots in response to ECM-compatible interactions reveals core gene regulons at early and mature developmental stages. (A) The barplot depicts the number of differentially expressed genes (DEGs) in poplar for each interaction at the early (grey) and mature (black) stages. Upregulated genes are represented in red and downregulated in blue. (B) Violin plot showing the results of Variance Partition Analysis of poplar genes. The variance of the genes that was mainly explained (≥ 0.5) by the fungal co-culture variable (above the blue dotted line) were selected for further analysis. (C) Upset plot representing the overlaps between the sets of DEGs for each poplar-fungal interaction and the selected genes highly variable according to the fungal co-culture. ECM core gene regulons at early and mature stages are highlighted in red. Pie charts represent the predicted topology composition of each regulon. Glob = Globular; SP = Signal peptide; TM = Transmembrane.

The “early ECM core gene regulon” was composed of 95 upregulated genes, with significant enriched molecular functions related to GO:0045735, nutrient reservoir activity; GO:0005506, iron ion binding; GO:0020037, heme binding; GO:0004497 monooxygenase; or GO:0016705 oxidoreductase activities, among others (**Table S5A**). Within these functions, a total of six genes presented nutrient reservoir activity annotation (**Table S5B**) and coded for small proteins (< 300 aa) with a signal peptide and a cupin domain with one to three germin-like motifs (**Fig. S2**). The nine significantly enriched genes presenting iron binding, heme binding and oxidoreductase and monooxygenase activity were all annotated as cytochrome P450 coding genes (**Table S5B**). No GO terms were significantly enriched in the 30 downregulated genes belonging to the “early ECM core gene regulon”.

Within the “mature ECM core gene regulon”, we found 61 upregulated genes, which were significantly enriched in GO:0009607, response to biotic stimulus and GO:0006468 protein phosphorylation biological processes (**Table S6A**). The two genes annotated as response to biotic stimulus were identified as pathogenesis-related coding genes from the pathogenesis-related bet v1 family. Up to nine upregulated genes were identified as protein kinases, with eight of them annotated as receptor-like kinases (RLKs) and one cytosolic calcium-dependant protein kinase. Among the RLKs, we found four cysteine-rich-stress-antifungal RLKs, three homologs of rust resistance kinase Lr10 from maize, and one G-type lectin S-RLK (**Table S6B**). On the other hand, we found 77 downregulated genes, which were mainly enriched in GO:0006979, response to oxidative stress (**Table S6A**), which corresponded with six downregulated peroxidase superfamily genes (**Table S6B**). We also found several GO terms related to cell wall remodelling (**Table S6A**). These genes were annotated as two xyloglucan endotransglucosylase / hydrolase 26, three plant invertase / pectin methylesterase inhibitors, two pectin-lyases and a glycosyl hydrolase 9C1 (**Table S6B**). Nine downregulated genes were annotated as pollen Ole e1 allergens (**Table 6B**) and eight of them presented a predicted signal peptide and three proline-rich extensin signatures apart from the pollen Ole e 1 domain (**Fig. S3**).

### Poplar ECM core gene regulon consists in general fungal-sensing and ECM-specific components

After the identification of poplar ECM core gene regulons at different developmental stages, we hypothesized that to be specifically regulated during ECM ontogenesis, they must be not regulated in response to non-compatible ECM fungal interactions. To test this hypothesis, we included in our analyses the RNAseq data from the non-compatible *P. tinctorius* and *S. citrinum* interactions at both developmental stages (**Fig. 1**). We identified a range of 774 and 1,768 DEGs (**Tables S7 and S8)** for each interaction and developmental stage (**Fig. 3A, Fig. S4**), representing a significant lower number of DEGs than ECM compatible interactions (**Fig. 3B**).

**Figure 3.**
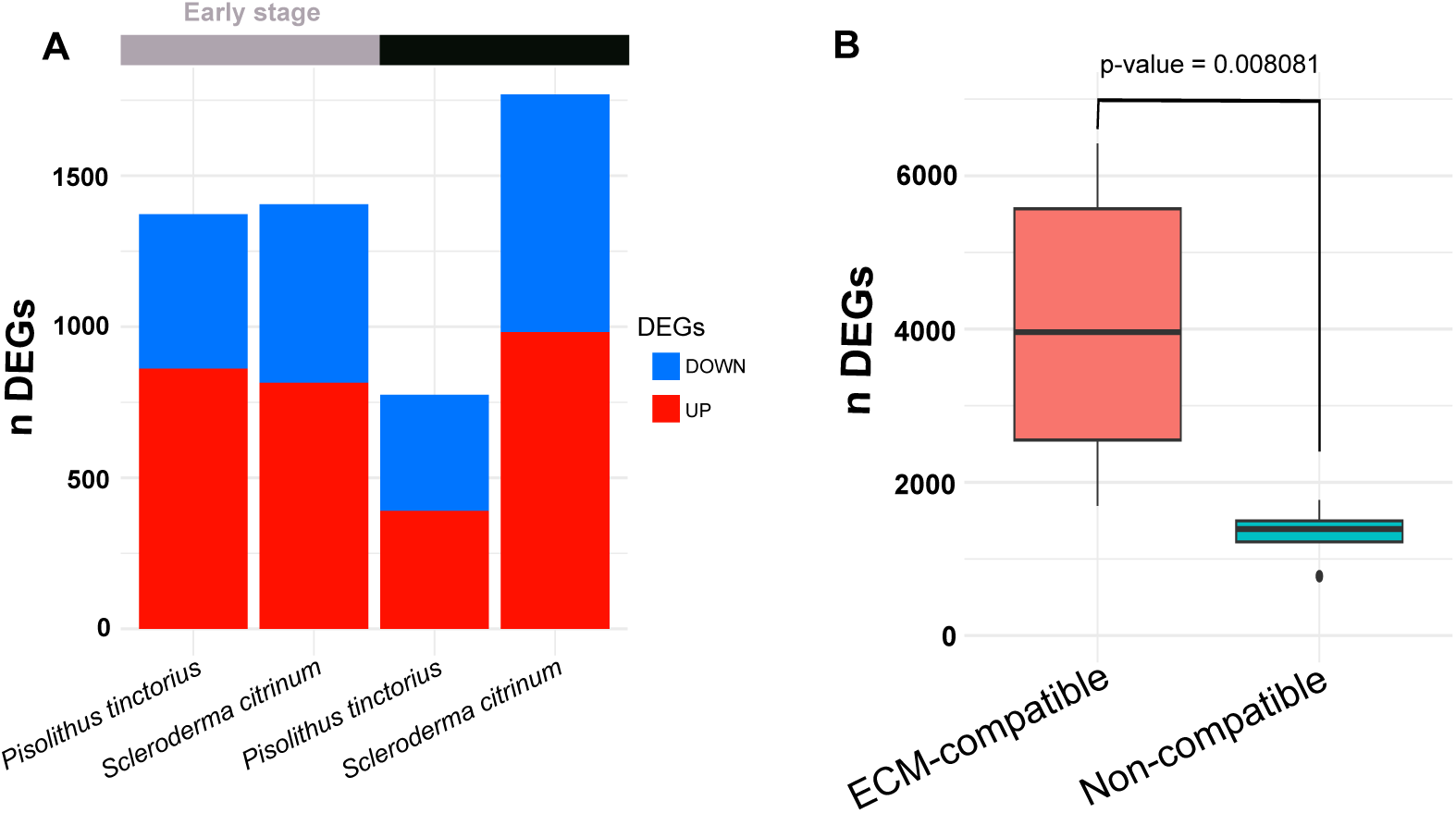
Interaction between poplar roots and non-compatible interactions cause weaker plant transcriptomic response than ECM-compatible interactions. (A) The barplot depicts the number of differentially expressed genes (DEGs) in poplar for each interaction at the early (grey) and mature (black) stages. Upregulated genes are represented in red and downregulated in blue. (B) Boxplot representing the number for poplar DEGs interacting with ECM-compatible and non-compatible fungal partners. Whiskers represent the limits of the 1.5 interquartile range. P-value refers to the Mann-Whitney-Wilcoxon test performed to decide whether the population distributions between ECM-compatible and non-compatible interactions were identical or not.

We overlapped the previously identified ECM core gene regulons with non-compatible DEGs. 51 out of the 125 genes (40.8%) from the early ECM core gene regulon were specific to ECM symbiosis i.e. not differentially regulated during non-compatible interactions (**Fig. 4A**). Three genes were downregulated, but the vast majority (48/51) of these ECM-specific genes among the “early ECM core gene regulon” were upregulated during ECM ontogenesis, with a subtilisin (Potri.014G026600) and two S-acyl fatty acid synthase thioesterase (Potri.001G177600 and Potri.001G177900) coding genes being the most upregulated genes. Eight genes were predicted to contain a signal peptide, including the mentioned subtilisin, a peroxidase and six small secreted proteins (SSPs), including two germin-like encoding genes, PADRE, PAR1, Derlin and an unknown gene (**Fig. 4B**). On the other hand, 20 out of the 125 genes (16.00 %) from the early ECM core gene regulon were commonly and in the same way regulated in all interactions and can be considered the general fungal-sensing part of the early ECM core gene regulon. Here, we found five different genes encoding for stress / antifungal receptor-like kinases, highlighting *Potri.017G040300*, which was the most upregulated gene in all interactions. We also found four out of the six secreted cupin encoding genes previously identified. Among the downregulated genes, we found a lipoxygenase homolog to *Arabidopsis thaliana LOX2* (**Fig. 4C**).

**Figure 4.**
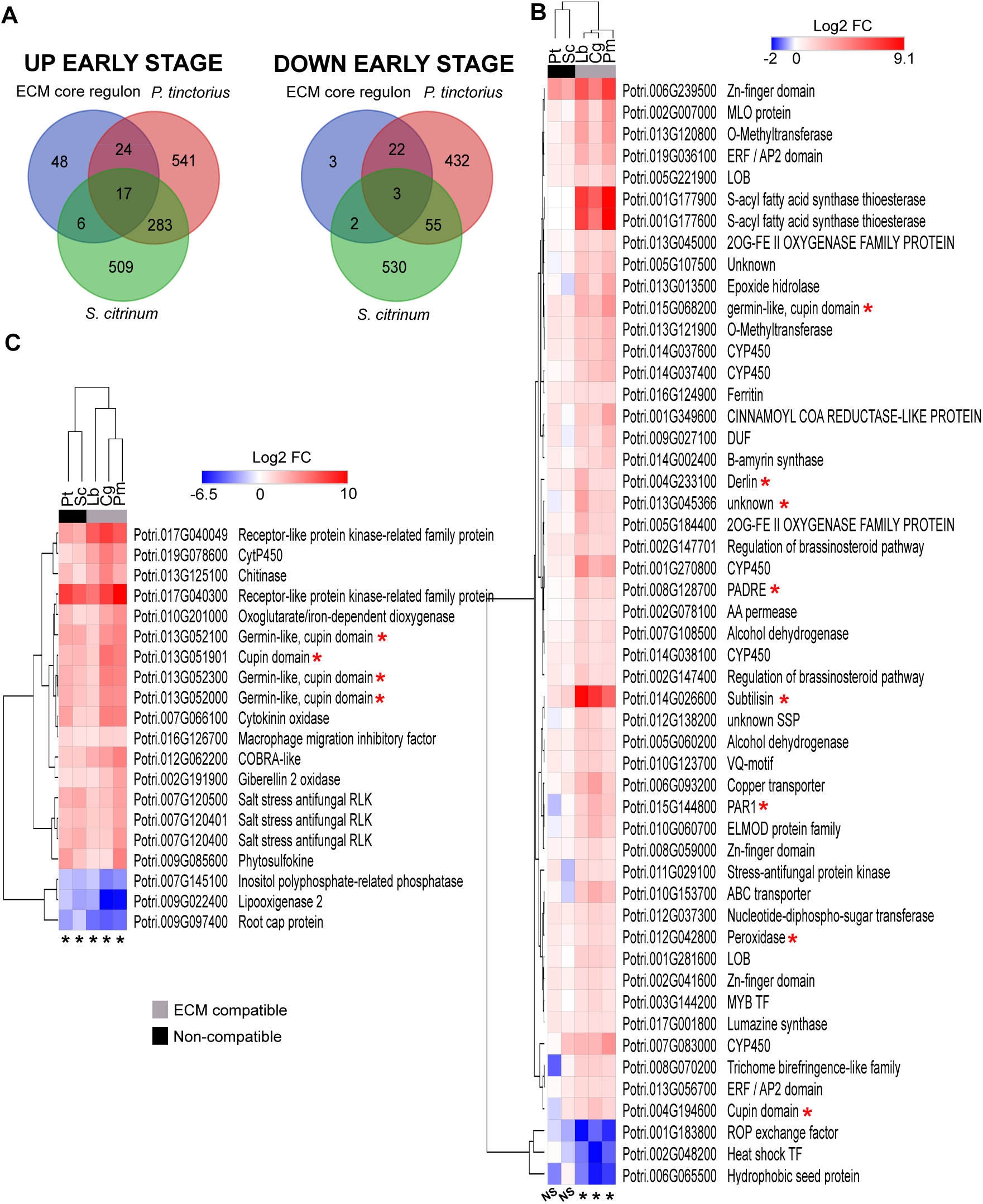
The early ECM core transcriptome of poplar consists of general fungal-sensing and ECM-specific genes. (A) Venn’s diagram depicting the gene overlaps between the previously defined “ECM core gene regulon” and differentially expressed genes in noncompatible interactions at early stage of ECM formation. (B) Heatmap showing the log2 Fold-Change of the 51 ECM-specific genes of the ECM core gene regulon at early stage. (C) Heatmap representing the log2 Fold-Change of 20 general fungal-sensing genes of the ECM core gene regulon at early stage. Significance according to the selected criteria (log2FC > 1; FDR < 0.01) is represented with a black asterisk (*) at the bottom of each heatmap. Although not significantly enriched according to these criteria, values for non-compatible interactions are also shown and represented with NS at the bottom of the heatmap. Red colour represents upregulation and blue colour downregulation in colonized vs uncolonized control roots for each poplar – fungus interaction. Red coloured asterisk represents upregulated (red) or downregulated (blue) genes encoding predicted secreted proteins.

Within the 138 mature ECM core gene regulon, 71 genes (51.45%) were specifically regulated during ECM development and not differentially regulated during non-compatible interactions (**Fig. 5A**). In contrast with ECM-specific genes at early time point, the vast majority (57/71) of these ECM-specific genes at mature stage were downregulated. 25 of them were predicted to be secreted proteins and most of them belonged to significantly enriched families in the mature ECM core gene regulon, such as peroxidases, pollen Ole e1 allergens and genes associated to plant cell wall degradation/ remodelling, targeting mainly pectin and xyloglucan **(Fig. 5B)**. The downregulation of two basic helix-loop-helix (bHLH, *Potri.001G294300* and *Potri.014G017100*) and an ethylene response factor / AP2 (*Potri.001G004700*) were also among the ECM-specific part of the mature ECM core gene regulon (**Fig. 5B**). We also found 14 upregulated genes, including three cysteine rich, stress antifungal RLKs (**Fig. 5B**). On the other hand, we identified 22 genes (17.65%) common to all interactions forming the general fungal-sensing component of the mature ECM core gene regulon. 21 of them were upregulated, highlighting four pathogenesis-related genes, several CYP450 and transporters (SWEET, ABC) (**Fig. 5C**)

**Figure 5.**
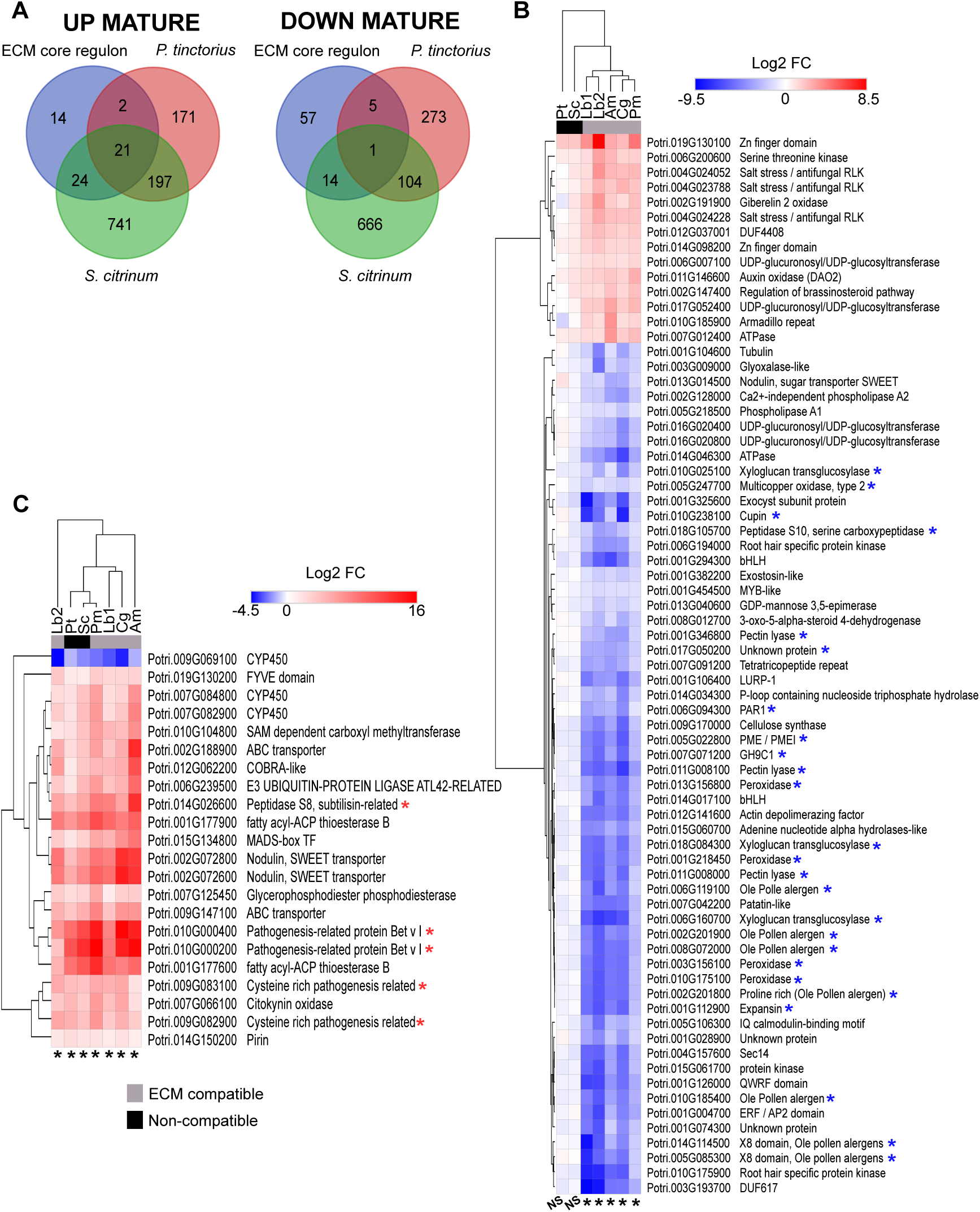
Poplar mature ECM core transcriptome is composed of general fungal-sensing and specific-ECM genes. (A) Venn’s diagram depicting the gene overlaps between the previously defined ECM core gene regulon and differentially expressed genes in noncompatible interactions at early stage of ECM formation. (B) Heatmap showing the log2 Fold-Change of the 71 ECM-specific genes of the ECM core gene regulon at mature stage. (C) Heatmap representing the log2 Fold-Change of 22 general fungal-sensing genes of the ECM core gene regulon at mature stage. Significance according to the selected criteria (log2FC > 1; FDR < 0.01) is represented with a black asterisk (*) at the bottom of each heatmap. Although not significantly enriched according to these criteria, values for non-compatible interactions are also shown and represented with NS at the bottom of the heatmap. Red colour represents upregulation and blue colour downregulation in colonized vs uncolonized control roots for each poplar – fungus interaction. Red coloured asterisk represents upregulated (red) or downregulated (blue) genes encoding predicted secreted proteins.

Finally, we also explored poplar genes that were commonly regulated in response to non-compatible interaction but were not present in the ECM core gene regulons (**Fig. 4A and 5A**). Both early and mature stages of non-compatible interactions displayed GO-enrichment related to pathogen-resistance functions **(Fig. 6A and B**), mainly GO:0009607, response to biotic stimulus; GO:0006952, defence response; GO:0009611 response to wounding; and GO:0004867, serine-type endopeptidase inhibitor activity. These terms mainly refer to secreted pathogen-related betv1 (PR-betv1) and serine endopeptidase inhibitor (SPI) encoding genes. Both gene families were not present in the ECM core gene regulon, except for two PR-betv1 genes found within the general fungal-sensing component of the mature ECM core (**Fig. 5C**). However, we found that, at early stage, all fungi elicited a strong poplar upregulation of both PR-betv1 and SPIs, except for *C. geophilum* (**Fig. 6C**). At mature stage, on the contrary, we found different trends between compatible and non-compatible interactions. Both non-compatible interactors still provoked a strong transcriptional defence response at mature stage, with only a decrease of 8.3 % for PRs and of 38.5 % for SPIs. The decrease on SPIs was mainly due to *P. tinctorius*, since it changed from twelve to four upregulated genes, whereas *S. citrinum* only shifted from fourteen to twelve (**Fig. 6D, Table S9**). On the other hand, ECM compatible interactors strongly decreased the activation of these defence-related gene families, especially in the case of SPIs. Only considering the interactions with available data for both early and mature stages, we found a decrease of 89.8 % in the number of upregulated SPIs and 66.7% for PRs (**Fig. 6D, Table S9**). On top of this, no downregulation of these families was found in non-compatible interactions at any stage. In ECM compatible interactions, during the transition from early to mature ECM, downregulation shifted from 0 to 8 for PRs and 1 to 11 for SPIs (**Fig. 6D**, **Table S9**).

**Figure 6.**
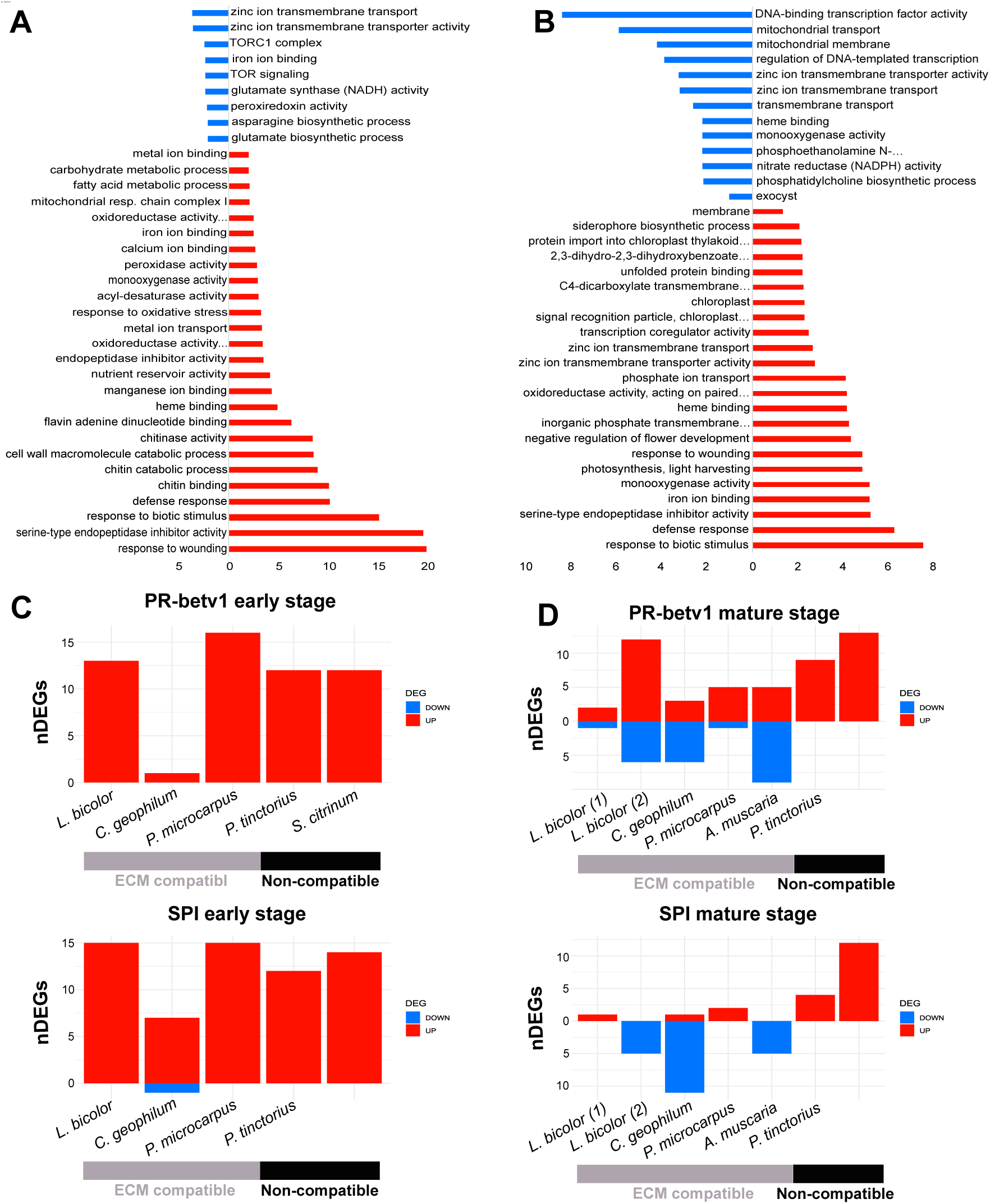
Differential regulation of plant secreted defence-related genes in ECM-compatible and non-compatible interactions. Significantly enriched Gene Ontology (GO) enriched terms in poplar non-compatible interactions at early (A) and mature (B) stages. Barplots depict the number of differentially expressed genes (DEGs) belonging to pathogenesis related betv1 (PR-betv1) and Serine endopeptidase inhibitors (SPI) gene families at early (C) and mature (D) stages.

### Poplar common symbiosis pathway does not belong to the ECM core gene regulon but is partially regulated during ECM symbiosis

Since poplar can establish both ECM and AM symbiosis and therefore possesses all genes of the Common Symbiosis Pathway (CSP), we explored their expression (**Fig. S5**) and regulation (**Fig. 7A**) in our datasets. We further included a selection of genes identified by Radhakrishan et al. (2020) as linked to AM and partially lost in non-AM plants (**Fig. 7B**).

**Figure 7.**
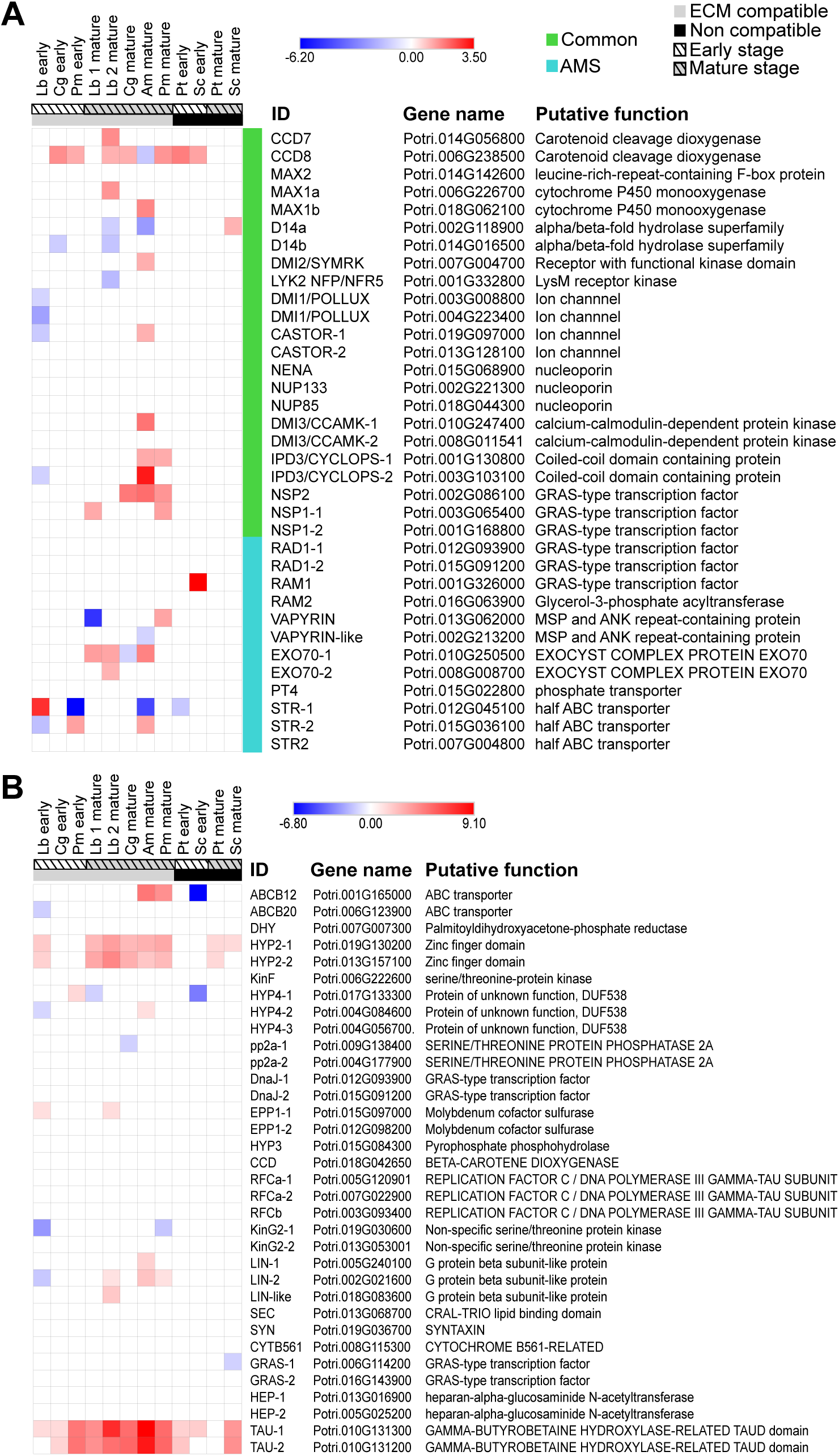
Genes involved in AM symbiosis are differentially expressed with poplar roots and ECM compatible and non-compatible interactions. (A) Common Symbiosis Pathway. Common, genes required for both symbioses with AM fungi and rhizobia; AMS, arbuscular mycorrhiza specific. (B) AM conserved genes identified by Radhakrishan et al. (2020).

CSP genes which in legumes are required for both symbioses with AM fungi and rhizobia **(Fig.7A**, common) are all expressed in poplar roots, independently from the colonization status or whether the interaction is compatible or non-compatible (**Fig. S5A**). From the downstream, AM symbiosis-specific genes **(Fig. 7A**, AMS), only *VAPYRIN* and *Exo70* genes involved in exocytosis, and *STR*, in transport, are expressed. For *RAD1*, *RAM1*, *RAM2*, *PT4* no or only background expression could be detected (**Fig. S5A**). Regarding the AM-conserved genes (**Fig. 7B**), no transcripts or only background level expression were detected for *ABCB12*, *DHY*, *KinF*, *Hyp4*-1, *pp2a*-2, *DnaJ*, *EPP1*-2, *HYP3*, *CCD*, *RFCa*, *KinG2*, *GRAS*-2 and *HEP*-2, while *ABCB20*, *SEC*, *SYN*, *CYTB561* and *GRAS-1* were constitutively expressed in all conditions (**Fig. S5B**).

Overall, the number of upregulated genes in compatible interactions was higher than in non-compatible interactions and include *MAX1* (with Lb, Am), *DMI2* (with Am), *CASTOR* (with Am), *DMI3* (with Am), *CYCLOPS* (with Am, Pm), *NSP2* (with Cg, Am, Pm), *NSP1* (with Lb, Pm), *VAPYRIN* (with Pm) and *Exo70* (with Lb, Cg, Am). *CCD8*, involved in strigolactone biosynthesis, was found up-regulated in both compatible and non-compatible interactions, while *RAM1* and *D14* were induced only in contact with the non-compatible *S. citrinum*. Overall, we found the regulation of 21 / 35 genes evaluated in ECM compatible interactions, while only 4 /35 were regulated in non-compatible interactions (**Fig. 7A**). Interestingly, genes coding for the Zinc-finger domain containing *HYP2* and the *GAMMA-BUTYROBETAINE HYDROXYLASE-RELATED TAU* were highly induced in all interactions (**Fig. 7B**).

### Sclerodermataceae fungi present differential gene expression patterns according to their ability to establish ECM interaction with poplar roots

To explore fungal transcriptional traits related to ECM compatibility, we also analysed the RNAseq data from three different Sclerodermataceae fungi in co-culture with *P. tremula x alba* at two developmental stages, early contact and mature stage. As stated above, *P. microcarpus* was the only Sclerodermataceae fungal partner that established a compatible ECM symbiosis with poplar (**Fig. 1**). We found a total of 1,168 DEGs (log2FC > |1|, FDR < 0.01) at early stage for *P. microcarpus* (**Table S10A**), a number strikingly higher than for *P. tinctorius* (235 DEGs, log2FC > |1|, FDR < 0.01, **Table S10B**) or *S. citrinum* (248 DEGs, log2FC > |1|, FDR < 0.01, **Table S10C**), which established non-compatible interactions (**Fig. 1 and Fig. 8A**). The KOG group “Information Storage and Processing” was overrepresented in the *P. microcarpus* upregulated genes at early stage compared to the non-compatible partners, with 117 DEGs *vs* 0 and 2 DEGs for *P. tinctorius* and *S. citrinum*, respectively (**Fig. 8A**). Among these 117 upregulated genes, we found 76 genes related to translation processes, mainly encoding for subunits of ribosomal proteins, 16 genes related with DNA replication or 11 related to chromatin structure (**Fig. 8B**). At mature stage, we found the opposite trend, since *P. microcarpus* was the fungi with the weaker transcriptional response (378 DEGs, log2FC > |1|, FDR < 0.01, **Table S11A**) compared to the non-compatible interactors *P. tinctorius* (566 DEGs, log2FC > |1|, FDR < 0.01, **Table S11B**) and especially *S. citrinum* (1,237 DEGs, log2FC > |1|, FDR < 0.01, **Table S11C**). Although the number of DEGs was different between fungi at mature stages, no striking differences were found in terms of KOG group composition between the datasets (**Fig. 8A**)..

**Figure 8.**
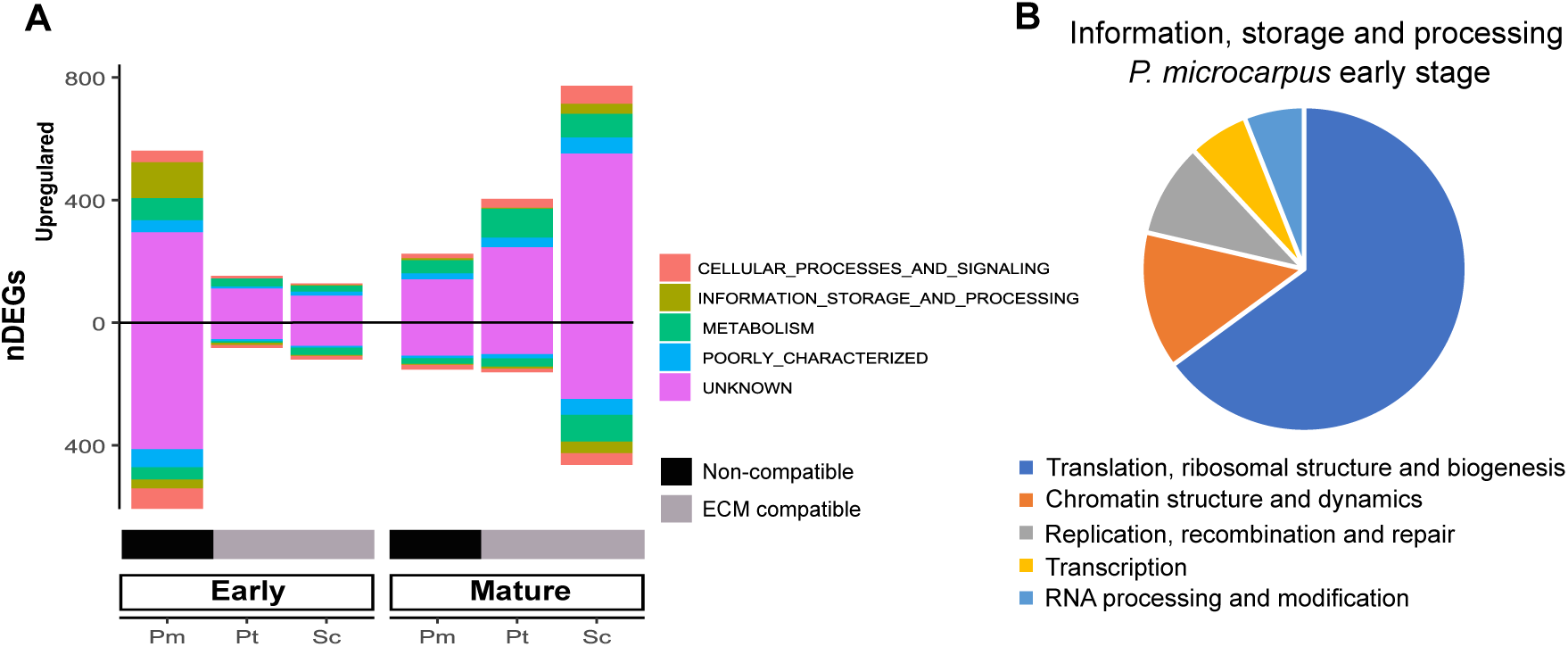
Gene expression patterns of Sclerodermataceae fungi in interaction with poplar roots reveal differences in genes related to information storage and processing at early stage. (A) The barplot depicts the number of differentially expressed genes from *P. microcarpus*, *P. tinctorius* and *S. citrinum* at early and mature stages of the interaction with *P. tremula x alba*. Their affiliation to KOG groups according to their *in silico* annotation is shown with different colours. Upregulated and downregulated genes are represented with positive and negative numbers, respectively. (B) The pie chart represents the KOG class distribution of Information, storage and processing genes from *P. microcarpus* at early stage.

Because SSP are prominent within the top 10 upregulated genes from each interaction (**Table 2**) and because fungal secretomes play a critical role for the establishment of ECM symbiosis (Kohler et al., 2015; Pellegrin et al., 2015; Miyauchi et al., 2020), we decided to compare the expression of the fungal secretome in the compatible and non-compatible interactions. At early stage, *P. microcarpus* regulated the expression of 141 putative secreted proteins. 102 of them were upregulated, and half of them (54) were small secreted proteins (SSPs) (**Fig. 9A**). Some of these SSPs encoding genes were among the top 10 most upregulated genes including *Pm683008*, *Pm688217*, *Pm4129416* or *Pm2071234* (**Table 2**). The *P. microcarpus* homolog (*Pm683339*) of the previously identified *PaMiSSP10b* (Plett et al., 2020) was also induced at this early stage of interaction. At mature stage, *P. micorcarpus* regulated a lower number of putative secreted proteins (**Fig. 9A**), but again most of them were upregulated and among them we found 20 SSPs encoding genes, including again *Pm683008* as one of the top 10 upregulated genes at mature stage (**Table 2**). *Pm683008* showed ≥ 85% similarity of sequence to *P. tinctorius Pt31842* and *Pt3134325*, and to *S. citrinum Sc27473* (**Fig. S5**). Interestingly *Sc27473* is also upregulated (log2FC 3.3) in the incompatible late stage (Table S11C). The three *P.tinctorius* TOP10 up-regulated SSPs (*Pt4353109, Pt4020897, Pt4013770*) are not the closest homologs of *Pm683008* but share some sequence homology, indicating a possible common origin. (**Table 2, Tables S10 and S11**). Moreover, the transcriptional response was stronger at mature than at early stage, contrary to the observed profile of expression of SSPs from the ECM compatible *P. microcarpus* (**Fig. 9A**). Additionally, we observed that hydrophobins were among the most upregulated genes for *P. tinctorius* at both early and late stages, and for *S. citrinum* at late stage (**Table 2**). Finally, because fungal Carbohydrates-Active Enzymes (CAZymes) are important for the development of mature ECM (Zhang et al., 2018; 2020), we studied the expression profile of all CAZYme families that were expressed in at least one condition and compared it with their homologs in the other fungal species. Overall, *P. microcarpus* was the fungus that upregulated more CAZYmes (**Fig. 9B**). Laccases (AA1), expansins (EXPN) or carboxyl esterase 4 (CE4)-encoding genes were commonly regulated by the three fungi, while GH5_15, CBM50, GH17, GH37 or GH28 and AA9 families were exclusively regulated in *P. microcarpus* i.e. during compatible interaction (**Fig. 9B**).

**Figure 9.**
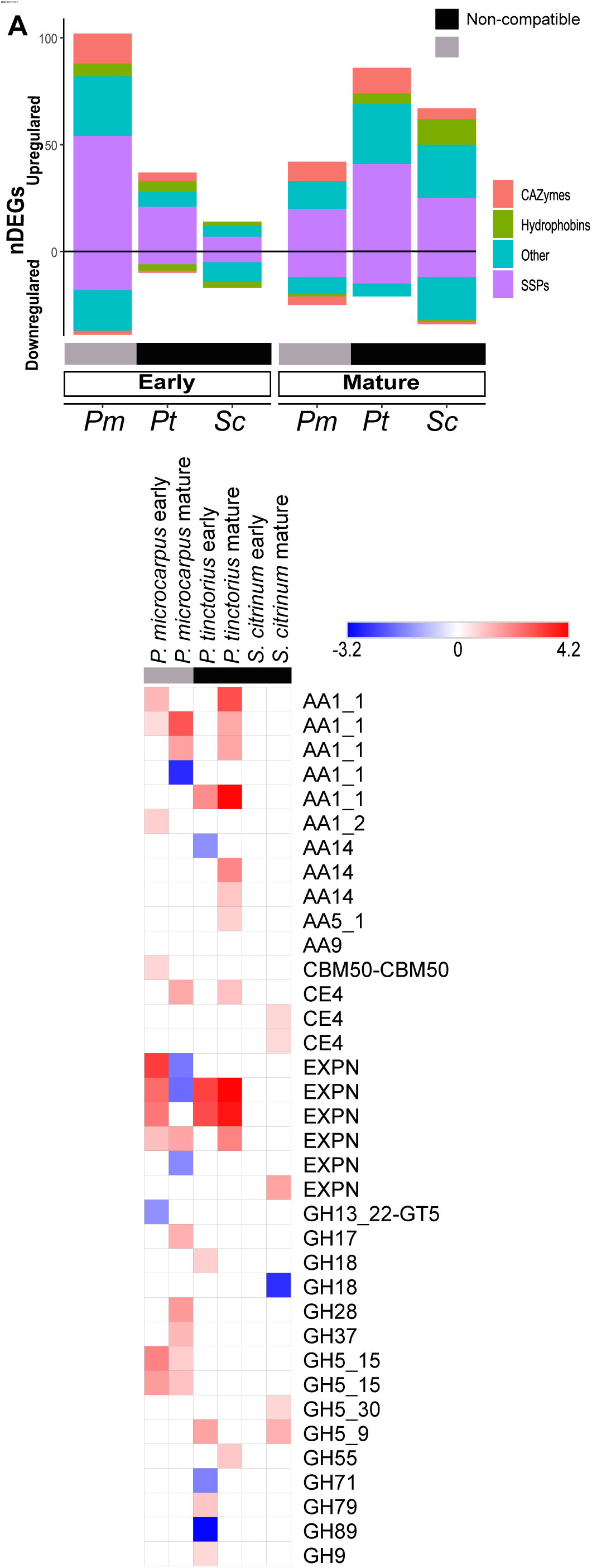
Secretomes of sclerodermataceae are differentially expressed according to ECM-compatbile or non-compatible interactions with poplar roots. (A) The barplot depicts the number of differentially expressed genes from *P. microcarpus*, *P. tinctorius* and *S. citrinum* at early and mature stages of the interaction with *P. tremula x alba*. Their annotation is shown with different colours. Upregulated genes are represented with positive numbers and downregulated genes with negative numbers. (B) The heatmap depicts the log2 fold-change of the expression of secreted CAZymes. Upregulated genes are represented in red and downregulated genes in blue. NR = Not regulated, in black. CAZyme-encoding genes are grouped according to their homology across species.

**Table 2.** Top 10 differentially expressed genes from Sclerodermataceae fungi at different developmental stages of the interaction with *P. tremula x alba*.

| *Pisolithus microcarpus* at early stage |  |  |  |  |  |
| --- | --- | --- | --- | --- | --- |
| Upregulated |  |  | Downregulated |  |  |
| Protein ID | Annotation | Log2FC | Protein ID | Annotation | Log2FC |
| 688217 | SSP | 5.31 | 4351830 | Activator of mitotic machinery Cdc14 phosphatase | -4.96 |
| 683008 | SSP | 4.68 | 4211579 | Hypothetical protein | -4.87 |
| 466641 | Zn(2)Cys(6) fungal-type DNA-binding | 4.58 | 4037734 | Anykrin repeat protein | -4.27 |
| 4129416 | SSP | 4.34 | 1774743 | Hypothetical protein | -3.27 |
| 4338441 | Symbiosis-regulated acidic polypeptide | 4.09 | 29985 | Zn-finger DNA binding | -3.06 |
| 686526 | Symbiosis-regulated acidic polypeptide | 3.91 | 3037960 | Hypothetical protein | -2.90 |
| 114059 | Hypothetical protein | 3.81 | 4297991 | Glutathione transferase | -2.84 |
| 4305455 | Amine oxidase | 3.81 | 448838 | Hypothetical protein | -2.51 |
| 440048 | Symbiosis-regulated acidic polypeptide | 3.78 | 4345076 | Mitochondrial ADP/ATP carrier | -2.49 |
| 2071234 | SSP | 3.75 | 4345466 | Hypothetical protein | -2.49 |
| *Pisolithus microcarpus* at mature stage |  |  |  |  |  |
| Upregulated |  |  | Downregulated |  |  |
| Protein ID | Annotation | Log2FC | Protein ID | Annotation | Log2FC |
| 4111547 | HSP20-like chaperone | 6.86 | 4229307 | Hypothetical protein | -3.72 |
| 572741 | HSP20-like chaperone | 5.72 | 4043478 | Heterokaryon incompatibly | -3.53 |
| 440048 | Symbiosis-regulated acidic polypeptide | 5.42 | 4369521 | Hypothetical protein | -3.08 |
| 4301408 | Major facilitator | 5.06 | 675652 | MYND finger | -2.64 |
| 6861 | CYP450 | 4.80 | 4327935 | Hypothetical protein | -2.63 |
| 6508 | Symbiosis-regulated acidic polypeptide | 4.27 | 4319684 | Laccase | -2.59 |
| 4577275 | Major facilitator | 4.16 | 4258660 | Hypothetical protein | -2.54 |
| 683008 | SSP | 4.16 | 12269 | Hypothetical protein | -2.51 |
| 4282329 | FMN binding | 3.95 | 690658 | SSP | -5.50 |
| 689604 | SSP | 3.89 | 4243894 | Hypothetical protein | -2.48 |
| ***Pisolithus tinctorius* at early stage** |  |  |  |  |  |
| **Upregulated** |  |  | **Downregulated** |  |  |
| **Protein ID** | **Annotation** | **Log2FC** | **Protein ID** | **Annotation** | **Log2FC** |
| 171326 | Hydrophobin | 5.44 | 20562 | Glutathione transferase | -7.33 |
| 891285 | SSP | 4.80 | 17316 | Sorbose reductase | -6.98 |
| 4111808 | Hypothetical protein | 4.64 | 920403 | Protein binding | -6.39 |
| 1001430 | Flavonol reductase/cinnamoyl-CoA reductase | 4.16 | 904946 | Zinc finger | -5.77 |
| 148654 | Hydrophobin | 3.98 | 4222525 | Hypothetical protein | -5.59 |
| 212195 | Hypothetical protein | 3.92 | 1005360 | Hypothetical protein | -5.48 |
| 1008833 | Hypothetical protein | 3.75 | 138292 | P-loop containing nucleoside triphosphate hydrolase | -5.37 |
| 4221927 | Hypothetical protein | 3.63 | 609947 | Hypothetical protein | -5.34 |
| 2628146 | Hypothetical protein | 3.37 | 21517 | Hypothetical protein | -5.12 |
| 3983026 | Hydrophobin | 3.34 | 891475 | SSP | -4.68 |
| ***Pisolithus tinctorius* at mature stage** |  |  |  |  |  |
| **Upregulated** |  |  | **Downregulated** |  |  |
| **Protein ID** | **Annotation** | **Log2FC** | **Protein ID** | **Annotation** | **Log2FC** |
| 212195 | Hypothetical protein | 6.90 | 4217061 | Hypothetical protein | -5.23 |
| 4353109 | SSP | 6.68 | 118669 | Hypothetical protein | -4.07 |
| 148654 | Hydrophobin | 6.05 | 9163 | 4-coumarate-CoA | -3.69 |
| 1003164 | Symbiosis-regulated acidic polypeptide | 5.86 | 1007633 | Hypothetical protein | -3.58 |
| 4020897 | SSP | 5.81 | 571198 | Hypothetical protein | -3.46 |
| 3983026 | Hydrophobin | 5.80 | 4224891 | glycerol-1-phosphatase | -3.40 |
| 4125539 | Hypothetical protein | 5.73 | 4129130 | Hypothetical protein | -3.30 |
| 171326 | Hydrophobin | 5.72 | 3358904 | Zn ion binding | -3.17 |
| 4013770 | SSP | 5.71 | 1001023 | Hypothetical protein | -3.00 |
| 1001308 | Hypothetical protein | 5.37 | 995413 | Hypothetical protein | -2.90 |
| ***Scleroderma citrinum* at early stage** |  |  |  |  |  |
| **Upregulated** |  |  | **Downregulated** |  |  |

| Protein ID | Annotation | Log2FC | Protein ID | Annotation | Log2FC |
| --- | --- | --- | --- | --- | --- |
| 142697 | CYP450 | 6.31 | 1170341 | Hypothetical protein | -8.06 |
| 10358 | Hypothetical protein | 4.63 | 1218978 | Hypothetical protein | -7.98 |
| 21074 | Methionyl aminopeptidase | 3.63 | 1018192 | Hypothetical protein | -4.06 |
| 1218425 | Hypothetical protein | 3.59 | 1222722 | HSP20-like chaperone | -2.75 |
| 1223812 | Hypothetical protein | 3.30 | 1210984 | Methyltransferase | -2.73 |
| 1223610 | SSP | 3.25 | 112173 | Isoprenylcysteine carboxyl methyltransferase | -2.62 |
| 1009768 | Hypothetical protein | 3.18 | 1073024 | Hypothetical protein | -2.51 |
| 49048 | Phosphatidylserine decarboxylase | 3.03 | 976814 | HSP20-like chaperone | -2.51 |
| 14443 | TPR repeat | 2.96 | 1211101 | Hypothetical protein | -2.46 |
| 1225039 | Hypothetical protein | 2.85 | 1211089 | Hypothetical protein | -2.36 |
| ***Scleroderma citrinum* at mature stage** |  |  |  |  |  |
| Upregulated |  |  | Downregulated |  |  |
| Protein ID | Annotation | Log2FC | Protein ID | Annotation | Log2FC |
| 1215971 | Hydrophobin | 6.60 | 1170341 | Hypothetical protein | -8.30 |
| 142697 | CYP450 | 5.77 | 119113 | Hypothetical protein | -4.47 |
| 1215978 | Hydrophobin | 5.38 | 976814 | HSP20-like chaperone | -4.28 |
| 1224799 | Hypothetical protein | 4.79 | 1211547 | Major facilitator superfamily | -4.18 |
| 1221589 | Hypothetical protein | 4.45 | 1217250 | Hypothetical protein | -3.92 |
| 1216465 | Ricin-B-related lectin | 4.36 | 1217249 | Hypothetical protein | -3.85 |
| 874652 | Hydrophobin | 4.20 | 1222722 | HSP20-like chaperone | -3.64 |
| 1210199 | Hypothetical protein | 4.15 | 349420 | Hypothetical protein | -3.53 |
| 1220978 | SSP | 3.96 | 8427 | Hypothetical protein | -3.51 |
| 1220549 | Hypothetical protein | 3.74 | 1217970 | Hypothetical protein | -3.51 |

## Discussion

Previous transcriptomic studies of both tree and fungi have aimed to decipher the two-way molecular dialogue allowing ectomycorrhiza ontogenesis. This research has predominantly focused on model systems such as *Populus sp.* – *Laccaria bicolor* (Martin et al., 2008; Veneault-Fourrey et al., 2014; Ruytinx et al., 2021), *Eucalyptus sp.*- *Pisolithus sp.* (Duplessis et al., 2005; Plett et al., 2015), or *Quercus sp.* – *Piloderma croceum* (Herrman and Buscot, 2007; Frettinger et al., 2007). However, only a limited number of studies (e.g. Liao et al., 2016) have interrogated at a genome-wide scale, the transcriptional responses of plants and ECM fungi to identify plant and fungi compatibility factors. Therefore, we profiled transcriptomes of poplar roots in contact with several compatible or incompatible ECM fungi to increase the knowledge on this topic. Our dataset demonstrates (1) the existence of plant genes that could act as “sensor” for fungal presence in the vicinity of the roots; (2) a minor, more general role of the common symbiosis signalling pathway in signal transduction; (3) inhibition of part of the plant immune response is a pre-requisite for compatible interactions to occur; (4) fungal CAZymes are the main actors of plant-cell wall remodelling leading *in planta* fungal growth and Hartig net development, and (5) the key role of fungal small secreted proteins as determinants of ECM compatibility.

### 1. The general fungal-sensing component of the ECM core gene regulon

A small portion of the identified ECM core gene regulon was also regulated in non-compatible associations, implying a function in general fungal recognition and response to changes in their surroundings rather than specific ECM functions. The intense upregulation of RLKs, germin-like and pathogenesis related genes during every fungal co-culture (**Fig. 4C; 5C**) aligns with this, since these plant gene families are well-known to respond to external biotic and abiotic factors (Morris and Walker, 2003; Liu and Ekramoddoullah, 2006; Dunwell et al., 2008). These poplar genes are thus good candidate for perception and response to fungal-associated molecular patterns in roots. However, other genes such as the SWEET transporters were less expected to belong to this group of genes. SWEET proteins are the main membrane transporters of sugars in plant cells and are known to be transcriptionally activated during different root mutualistic symbioses (Kryvoruchko et al., 2016; Manck-Götzenberger and Requena, 2016). Additionally, PtaSWEET1c plays a crucial role facilitating glucose and sucrose transport at the symbiotic interface of *Populus* - *L. bicolor*. *PtaSWEET1c* is transcriptionally activated in response of *L. bicolor* co-culture even without physical contact (only diffusible signals were shared), but the activation is only maintained over time when direct contact occurred (Li et al., 2024). Therefore, we anticipated that *SWEET* gene(s) would be detected as members of the ECM-specific component. However, we found two closely related homologs of *PtaSWEET1c* (Potri.002G072700): *PtaSWEET1a* (Potri.002G072600) and *PtaSWEET1b* (Potri.002G072800), highly upregulated in response to all fungal co-cultures, regardless the compatibility (**Fig. 5C**). The baker yeast uses Snf3 and Rgt2 transporters as glucose sensors that generate a signal for induction of expression of genes encoding hexose transporters (Ozcan et al.,1998). We propose the hypothesis that these poplar PtaSWEET1a and PtaSWEET1b proteins act as low or high affinity sugar sensors (as the yeast RGT2 and SFN3) to generate signal for further induction of other SWEET transporter (such as PtaSWEET1c) required for functional symbiosis. In addition, SWEET sugar transporters play a role in the distribution of sugars along the root, which sustain spatial colonization of the root-associated bacteria (Loo et al., 2024). We can thus speculate of a role in spatial root colonization of fungi. Deeper functional studies should be performed to prove these hypotheses and to provide insights into the biological implications and spatio-temporal dynamics of these transporters.

### 2. The role of common symbiosis pathway in mycorrhizal compatibility

Mycorrhizal symbiosis is an ancestral plant trait imprinted in plant genomes as a signalling pathway called the Common Symbiosis Pathway (CSP) (Olroyd, 2013; Genre et al., 2020). This pathway is conserved in all plant lineages capable of establishing intracellular-forming symbioses, including root nodule symbiosis and arbuscular, orchid or ericoid mycorrhiza (Radhakrishnan et al., 2020). In poplar, which can establish both AM and ECM symbioses, important components of the CSP, such as *CASTOR/POLLUX* and a calcium/calmodulin-dependent protein kinase (*CCaMK*) genes, are involved in the interaction with the ECM fungus *Laccaria bicolor* (Cope et al., 2019). This pathway is absent in gymnosperm lineages exclusively forming ECM symbiosis, such as *Pinus* spp. (García et al., 2015; Bravo et al., 2016), but partially present in ECM-only species within the Fagales clade (Li et al., 2024). Additionally, certain CSP genes are upregulated in ECM roots of *Castanea mollissima*, including the *CASTOR* and *DMI3/CCaMK* genes known to be involved in ECM symbiosis in poplar (Li et al., 2024). Our results show that the transcription of a subset of CSP genes in poplar, including the aforementioned *CASTOR* and *DMI3/CCaMK*, is constitutively expressed in poplar roots. Overall, our results indicate that plants are constitutively ready to use this pathway to initiate symbiotic interactions and that the transcriptional activation of specific subsets of this pathway is associated to mycorrhizal compatibility and the establishment of ECM symbiosis. The exact mechanism by which these genes enable ECM ontogenesis and how ECM-exclusive plant species that do not present these genes in their genomes are also capable of forming ECM symbiosis needs to be explored in the future.

### 3. Growth-defence trade-off responses are common to all interactions but modulation of JA-related plant immunity at mature stage is specific to ECM-compatible interactions

According to previous reports, *L. bicolor* interaction with poplar roots causes the alteration of diverse plant hormonal pathways (Basso et al., 2020). In the context of the growth-defence trade-offs, gibberellins (GA), cytokinin (CK), brassinosteroids (BR) and auxin are associated to plant growth, while jasmonate (JA), ethylene (ET) and salicylic acid (SA), are associated with defence (He et al., 2022). We found upregulation of a gibberellin-2-oxidase and a cytokinin oxidase in all plant-fungal interactions (**Fig. 4C; 5C**). Two BR regulators at early and mature stages and an auxin oxidase at mature stage were only significantly upregulated in the case of compatible interactions, but in both cases a trend towards upregulation can be also observed for non-compatible interactions (**Fig. 4B; 5B**). The gibberellin, auxin or cytokinin oxidases are involved in the inactivation of their respective hormone bioactive forms (Thomas et al., 1999; Jones and Schreiber 1997; Zhang and Peer, 2017). BR regulators control BR levels *in planta* (Yuan et al., 2007). Overall, the transcriptional activation of growth-related hormone catabolism suggests a repression of plant growth-responses, which occurs in all type of interactions but is more pronounced in ECM-compatible interactions than in non-compatible ones. Concomitantly, we also observed upregulation of pathogenesis related (PR) and serine proteinase inhibitors (SPI) genes at early stages of both compatible and incompatible interactions (**Fig. 6C**). PR genes are associated to SA signalling (White, 1979) and Systemic Acquired Resistance (SAR) (Durrant and Dong, 2004), while proteinase inhibitors are linked to JA signalling (Farmer et al., 1992). Therefore, our data provide transcriptional evidence of growth-defence crosstalk in early stages of root-beneficial fungi interactions, regardless of compatibility status, favouring JA- and SA-related defence over GA-, BR-, CK- and auxin-mediated growth. Together with the up-regulation of poplar receptor-like kinase at early time-point (se point 1 of the discussion), our data identify the key transcriptionally activated players likely involved in how roots perceive fungi.

Following the early activation of plant defences, we also highlighted down-regulation of plant defence in later time-point (Fig.5). For example, the homolog of *COBRA-like 4* gene in *A. thaliana*, *Potri.012G062200* is upregulated at both stages of all type of interactions (Fig. 4C, 5C). This gene is involved in cellulose deposition in secondary cell walls (Sato et al., 2010; Xue et al., 2024), negatively interacts with JA pathway in *A. thaliana* (Ko et al., 2006) and is associated with Hartig net depletion in *MYC2* overexpressing poplar mutants (Marqués-Gálvez et al., 2024). Within the specific component of the ECM core gene regulon, we found downregulation of several transcription factors, such as bHLHs and ERFs (**Fig. 5B**), which are typically associated to JA-related defences (Grennan, 2008; Goossens et al., 2017). ECM-specific downregulation in the transition from early to mature stage was stronger for the JA-associated SPIs than for the SA-related PR genes (**Fig. 6D**, **Table S9**). This concurs with the crosstalk mechanism described in poplar - *L. bicolor* interactions, where *L. bicolor,* through the secretion of the fungal effector MiSSP7, regulates the activation of the MYC2 master defence factor. This regulation prevents the activation of JA-related defences, including other bHLHs and ERF transcription factors, terpenes, chitinases and proteinase inhibitors (Plett et al., 2011; 2014; Daguerre et al., 2020; Marqués-Gálvez et al., 2024). While dampening JA-responses seem a common trait in late ECM, *MiSSP7*, like many fungal effectors, is an orphan gene with low sequence homology (Kohler et al., 2015). The specific mechanisms by which different fungal clades, with their distinct effector toolkits, achieve to the same outcome (e.g. local modulation of JA-related defence) need to be thoroughly examined. Modulation of the defence response at mature stage of ECM-compatible interactions is more pronounced for JA-related than for SA-related defences. To the best of our knowledge, there is no report of fungal effectors targeting endogenous SA signalling in ECM symbioses, in contrast with what is known for plant pathogenic fungi (Tanaka et al., 2015). The partial activation of SA-related genes in mature ECM-compatible interactions could be related to primary defence against the symbiotroph but also to putative SAR, which has been described for ECM (Dreisschhoff et al., 2020). In conclusion, our data suggests that impeding SA and JA-mediated responses is a common feature for the establishment of compatible ECM, with less inhibition of SA-mediated responses compared to the highly impaired JA-mediated responses.

### 4. Are fungal CAZymes taking over plant cell wall remodelling?

Plant cell wall is the place of the earliest communication event between plants and ECM fungi and CAZymes are the most important enzymes involved in fungal and plant cell remodelling during the establishment of Hartig net (Veneault-Fourrey et al., 2014). Down-regulation of plant cell wall remodelling machinery of the host appeared as a common feature in ECM-compatible interactions. (**Fig. 5B**). Plant proteins targeting pectic polysaccharides such as pectin lyases (Yadav et al., 2009), xyloglucan endotransglucosylase/hydrolases (Van Sandt et al., 2007), plant invertases / pectin methylesterase inhibitors (Coculo and Lionetti, 2022), cellulose and hemicellulose were highlighted. Several homologs of the pollen Ole e1 allergen were also massively downregulated (**Fig. 5B**). These proteins are known to play a role in cell wall remodelling in the context of the pollen tube (Alché et al., 2004), and some of them are highly expressed in *Arabidopsis* roots (Hu et al., 2014). Plant peroxidases can be associated to cell wall loosening and stiffening (Francoz et al., 2015), and their downregulation (**Fig. 5B**) may also be related to plant cell wall remodelling machinery downregulation.

Concomitant to this down regulation of the host CAZYmes gene repertoire, our data highlighted upregulation of key fungal CAZymes specifically to ECM symbiosis. This might seem can be counterintuitive because the reduction of fungal CAZyme sets in the genomes of ectomycorrhizal fungal is a major trait of their convergent evolution towards ECM symbiosis from their saprotrophic ancestors (Kohler et al., 2015). Despite this massive loss, fungal glucanase (GH5) and endopolygalacturonase (GH28) still play a significant role for these mutualistic fungi. For example, in *P. tremula x alba* x *L. bicolor* ECM, a β-1,4 endoglucanase GH5 is located at the periphery of hyphae forming the Hartig net and the mantle, and its silencing results in decreased HN formation (Zhang et al., 2018). *L. bicolor* LbGH28 is upregulated during ECM symbsiosis and encodes an endopolygalaturase with its highest activity towards pectin and polygalacturonic acid. It is located in both fungal and plant cell walls at the symbiotic hyphal front and RNAi mutants have a lower ability to establish ECM (Zhang et al., 2022). A pectin methylesterase of *L. bicolor* (*LbPME1*) has also been characterized to play a role in ECM formation potentially through homogalacturonan de-esterification (Chowdhury et al., 2022). We found glycosyl hydrolase (GH) 5_15, 28 and 37 families exclusively upregulated in *P. microcarpus* ECM-compatible fungal partner, suggesting a role for these genes in HN formation and, therefore, in ECM compatibility (**Fig. 9B**). Overall, our data provide another line of evidence of the crucial role that fungal GH5_15 and GH28 transcriptional activation play during ECM ontogenesis. Furthermore, this mechanism seems not to be singular of specific interactions, but general to different plant-fungal combinations. These genes are not only key to intercellular colonization but also strong candidates to be compatibility factors in ECM interactions.

Intriguingly, at early stages of the interaction, when there is no fungal colonization of root cell intercellular spaces, significant upregulation of fungal CAZymes were found. The upregulation of laccases (AA1) and expansin-like proteins (EXPN) is a shared trait of *P. microcarpus* and *P. tinctorius*, that can establish a mantle. However, this was not observed with *S. citrinum*, which did not interact with poplar roots at all (**Fig. 1 and 9**). Veneault-Fourrey et al. (2014) found *L. bicolor Exp1* protein to be upregulated during ECM ontogenesis and located in fungal cell walls within ECM structures. Our data supports this claim also for Sclerodermataceae fungi, since EXPN upregulation during early stages suggest a role in hyphal aggregation and root surface attachment for mantle formation during the first stages of ECM establishment. Their role in hyphal colonization of root intercellular spaces is more doubtful: whereas the non-compatible *P. tinctorius* maintains upregulation at mature stage, the ECM-compatible *P. microcarpus* deactivates and even downregulates the majority of their *EXPN* set **(Fig. 9B**). In summary, our data supports the important role of EXPN-like proteins at early stages of interaction as a general trait of ECM symbiosis, but their spatio-temporal dynamics must be resolved to reach further conclusions about their role for *in planta* colonization.

Overall, the crosstalk of fungal up- and plant down-regulation of CAZymes and related genes suggest that during ECM symbiosis, fungi take the control over the host machinery for plant cell plant remodelling. Our data suggest that the ability to control the host cell wall remodelling and “make room” for colonization could be a determinant of ECM compatibility. We could hypothesize that fungi are able to control endogenous plant cell wall remodelling machinery thanks to unknown signalling pathways, so that they can take control over this process.

### 5. Fungal small secreted proteins are key determinants of ECM compatibility

In the past years, there has been an effort to characterize fungal MiSSPs that could act as fungal effectors of mycorrhizal symbiosis (Kloppholz et al., 2011; Plett et al., 2011, 2014; Sędzielewska and Brachmann, 2016; Voß et al., 2018; de Freitas et al., 2018; Kang et al., 2020), similarly to what occurs in pathogenic plant-fungal interactions. The three Sclerodermataceae transcriptomes generated *de novo* for this study presented upregulation of SSPs at every stage studied. *P. microcarpus* was the only one establishing a compatible ECM interaction (**Table 1**, **Fig. 1**) and presented the greatest number of upregulated SSPs (**Fig. 9**), which were especially abundant within the top 10 upregulated genes (**Table 2**). This increased expression of SSPs could be related to enhanced crosstalk and therefore compatibility between partners. *Pm683008* was the only SSP continuously upregulated at early and mature stage of ECM interaction (**Table 2**). Surprisingly, the closely related *Sc27473* (**Fig. S6**) was also induced at late stage of incompatible interaction and three *P. tinctorius* SSPs showing some sequence homology with *Pm683008* were among the TOP10 induced transcripts at late stage. According to this, *Pm683008* and the related SSPs from the incompatible interactions could be good candidates to study mycorrhizal compatibility between Sclerodermataceae fungi and poplar roots. In addition, *Pm683008* was also identified by Plett et al., (2021) as a core SSP in different strains of *P. microcarpus* in compatible ECM interaction with *Eucalyptus grandis*. Host-specificity factors have been much more studied in pathogenic than in mutualistic plant fungal interactions. Plant pathogenic fungal specificity is determined by fungal effectors and the presence of host resistance and susceptibility proteins (Li et al., 2020). Therefore, our data also indicates that *Pm683008* is a candidate to play a role in determining *P. microcarpus* host-specificity range. Future efforts aiming to functionally characterize this interesting fungal effector and putative susceptibility or resistance proteins in its hosts should be performed to prove this.

## Conclusions

In this work, we present an extensive study of plant and fungal transcriptomes to find compatibility factors that determine the outcome of ECM symbiosis. We found certain traits that were previously known to be important for singular ECM symbiosis - mainly studied in *Populus* spp. x *L. bicolor* model system - and discussed their implications in host-partner compatibility. The secretion of fungal-specific effectors, the modulation of plant defence pathways and the fungal control over plant apoplastic remodelling through key fungal glycosyl hydrolases, along with the coordinated regulation of plant and fungal cell wall remodelling enzymes, appear to be the key factors determining ECM compatibility. We also highlight the possible role of certain components of the Common Symbiosis Pathway in ECM symbiosis, which has been understudied so far compared to other root mutualistic symbioses. Although other transcriptional traits have not been discussed here, our transcriptome dataset should serve as a strong foundation for hypothesis-driven functional studies of plant and fungal genes that are crucial to study plant-fungus compatibility in ectomycorrhiza symbiosis.

## Materials and Methods

### Experimental design

In this work we analysed poplar and fungal transcriptomes in response to compatible and non-compatible ECM interactions. We used the ability of *in planta* colonization by establishing the Hartig Net as a proxy for ECM compatibility (Barker et al. 1998). Seven poplar - fungus datasets, in five independent experiments and six different fungal partners were included in the analysis (**Table S1**). When available, we analysed transcriptomes at two different stages of ECM formation: early and mature. The early stage of ECM establishment was defined as when roots and fungi have entered contact forming the mantle, but no *in planta* colonization was observed. Mature stage was defined as when *in planta* colonization forming HN was observed. Phenotypical characterization to ascertain ECM compatibility and the stage of the interaction was determined in previous works for *L. bicolor* S238N, *C. geophilum* 1.58 and *A. muscaria* MEII interactions (Basso et al., 2020, Ruytinx et al., 2021, de Freitas Pereira et al. 2018, Kohler et al., 2015), while it was characterized *de novo* for Sclerodermataceae fungi (*P. microcarpus*, *P. tinctorius* and *S. citrinum*) in this work (see next section). Fungal transcriptomes originating from previous experiments have been already analysed and thus, they were not included in our analyses. Only *de novo* generated fungal transcriptomes (Sclerodermataceae) were included in this work. Plant transcriptomes have never been analysed nor published, except for one *P. tremula x alba – L. bicolor* interaction (Basso et al., 2020). To be able to compare between interactions, all poplar transcriptomes at the different stages available were included in this study (**Table S1**).

### *Populus tremula x alba* and *Sclerodermataceae* in vitro co-culture

*Pisolithus microcarpus* 441, *Pisolithus tinctorius* Marx 270, and *Scleroderma citrinum* Foug A, fungal strains belonging to the Tree-Microbe Interactions Department, INRAE Grand-Est Nancy, France, were utilized in this work. Fungal cultures were kept on Modified Melin-Norkans (MMN) culture medium (Marx 1969) at 28 °C and transferred to new culture medium every 20 days. The hybrid poplar *Populus tremula* x *Populus alba* 717-1B4 INRAE clone were grown on half-strength Murashige and Skoog (MS) medium (Murashige and Skoog, 1962) in glass culture tubes under a 16-h photoperiod at 25°C in a growth chamber. They were micropropagated in vitro every 3-6 months until use for co-culture. In vitro interaction was established between *P. tremula* x *alba* and *P. microcarpus*, *P. tinctorius* and *S. citrinum* following the methodology described by Felten et al., (2009). Briefly, plants were micropropagated *in vitro* in Murashige and Skoog (MS) medium (Murashige and Skoog, 1962), with indole-butiric-acid (IBA) to synchronize rhizogenesis. In parallel, the MNM medium with sugar, phosphorus and nitrogen (Brun et al.,1995) was lined with cellophane membranes. Ten to twelve agar disks containing fungal mycelium from each fungal strain were placed on the cellophane membranes and kept at 25 °C for 20 days. Following this, two hybrid poplar plants/dish were transferred onto the fungal mycelium. Six plates, for a total of n = 12 biological replicates for each interaction were included in the experiment. Petri dishes were incubated in a growth chamber at 25 °C with 16 h light.day-1 for 30 days. A control treatment without fungal inoculation was also included.

Every seven days for *P. microcarpus* and *P. trinctorius*, and only at 7- and 28-days post inoculation (dpi) for *S. citrinum*, the petri dishes were scanned by Epson V700 Photo Perfect to follow root development. The number of colonized and non-colonized lateral root tips was measured using ImageJ analysis (http://imagej.nih.gov/ij/docs/index.html) and the image analysis software SmartRoot (Lobet, & Draye 2013).

Root tips presenting or not ECM morphotypes were sampled at 7 and 28 dpi, based in two different timestamps in ECM development in vitro: Pre-symbiosis (first contact and mantle formation, 7 dpi) and colonization (Hartig net (HN) establishment, 28 dpi). Sampled root tips were dehydrated in an ethanol series (10, 30 and 50% v/v) for 1 h and fixed in 70% ethanol (v/v) for 24 h at 4°C. The samples were incubated in 80%, 95% and absolute ethanol for 1 hour before infiltration. The infiltration was carried in Technovit 7100 resin (Kylzer and Co. GmbH, D-6393 Wehrheim/Ts) using increasing mixtures of ethanol and Technovit 7100 resin with hardener I (benzyl peroxide) for 2 h. The samples were transferred to moulds with 100% inclusion solution (Technovit 7100 resin with hardener II) for 2-4 h. Polymerization was carried out at room temperature. Cross sections (5 µm) were cut with a MICROM HM 355S rotary microtome and stained using toluidine blue (0.05%). The cross sections were then examined with an inverted microscope (ZEISS Axio Observer Z.1) equipped with a digital camera at suitable magnification. Photographs were taken by the digital camera and analysed using software Axiovision release 4.8.2 (06.2010) (Carl Zeiss). The presence or absence of mantle and HN were evaluated. A minimum of four replicates was used for statistical analysis.

### RNA sequencing and poplar transcriptome analysis

As stated above, RNAseq datasets of poplar interaction with *L. bicolor*, *C. geophillum* and *A. muscaria* were previously generated, and were retrieved from Short Read Archive (SRA) database (https://www.ncbi.nlm.nih.gov/sra). RNA extraction and sequencing details for these datasets can be consulted in Basso et al., (2020), Ruytnix et al., (2021), de Freitas Pereira et al. 2018 and Kohler et al., (2015). RNAseq dataset of poplar interacting with *P. microcarpus*, *P. tinctorius* and *S. citrinum* were generated *de novo* for this study (**Table S1**). Total RNA from root tips at 7- and 28-dpi was extracted using the CTAB-lithium chloride (LiCl) method (Chang et al., 1993). Assays for the quantification and integrity check were conducted using an Experion Automated Electrophoresis Station (Bio-Rad, Hercules, CA, USA). Preparation of libraries and 2 x 100bp Illumina HiSeq mRNA sequencing (RNA-Seq) was performed by Transcriptome platform of GenoToul (INRA Toulouse, France). After sequencing, raw reads from both previous and *de novo* generated datasets were analysed using FASTQC for quality control (Andrews et al., 2010), paired, trimmed, and mapped against *Populus trichocarpa* v4.1 transcripts (https://phytozome-next.jgi.doe.gov/info/Ptrichocarpa_v4_1) using CLC Genomics Workbench v21. Reads that mapped in genes that contained in average less than 5 raw mapped reads across all the conditions were removed.

For all interactions, before performing differential expression, quality assessment was performed by exploring log2 normalized counts, plotting principal component analysis and performing hierarchical clustering. After removal of inconsistent replicates, the RNA-seq data were adequate for further analyses (**Fig. S7, S8**). Differential transcription levels of the reads were calculated with DESeq2 (v1.44.0, Love et al., 2014). It was performed for each plant-fungal interaction comparing inoculated vs non-inoculated plants at two different developmental stages, when possible (**Table S1**). Condition-specific differentially expressed genes (DEGs) with a false discovery rate (FDR) < 0.01 and a log2 fold-change (Log2FC) ≥ |1|, were identified for each pairwise condition. Volcano plots were used to illustrate and highlight DEGs.

Mapped reads were also normalized using Variance Stabilizing Transformation (vst) function from DESeq2 and Variance Partition Analysis (VPA) was performed (v1.34.0, Hoffman and Schadt, 2016). For VPA the following formula was defined:

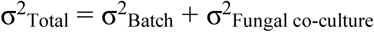

where σ^2^_Batch_ is the variance associated to the technical differences among all the experiments included in this analysis and σ^2^_Fungal co-culture_ is the variance associated to the presence or absence of fungi in co-culture. To identify the expression of a gene as mainly driven by a certain variable (σ^2^_x_), the variance associated to σ^2^_x_ must be ≥ 0.5. We overlapped the results from differential expression analysis and VPA to using UpsetR (v1.4.0, Conway et al., 2017) and Venn’s diagrams (https://bioinformatics.psb.ugent.be/webtools/Venn/). Additionally, TopGO (v2.42.0) R package was used for Gene Ontology (GO) enrichment analysis (Alexa and Rahnenfuhrer, 2022), with fisher test as statistical method and a p value threshold ≤ 0.01. To annotate specific groups of genes, their sequences were scanned using InterProScan (https://www.ebi.ac.uk/interpro) to find conserved domains and analysed using DeepTMHMM (Hallgren et al., 2022) to reveal protein topologies and possible signal peptides and transmembrane domains.

### RNA sequencing and *Sclerodermataceae* fungal transcriptome analysis

RNA extraction and sequencing details for *P. microcarpus*, *P. tinctorius* and *S. citrinum* free living mycelium and in co-culture with poplar at 7- and 28-dpi were as explained in the previous section. Raw reads were analysed using FASTQC for quality control (Andrews et al., 2010), paired, trimmed, and mapped against *P. microcarpus* (https://mycocosm.jgi.doe.gov/Pismi2/Pismi2.home.html), *P. tinctorius* (https://mycocosm.jgi.doe.gov/Pisti2/Pisti2.home.html) and *S. citrinum* (https://mycocosm.jgi.doe.gov/Sclci1/Sclci1.home.html) genomes. Quality control of RNAseq data, filtering and differential expression analysis between colonized and free-living mycelium controls were performed as for poplar datasets. Special focus on fungal secretomes focusing on Carbohydrate-Active enzymes (CAZYmes), lipases, proteases and small secreted proteins was performed due to their reported importance in the establishment of ECM symbiosis (Kohler et al., 2015; Pellegrin et al., 2015; Miyauchi et al., 2021). The RNA-seq data were adequate for further analyses (**Figs. S9, S10 and S11**). Differential transcription levels of the reads were calculated with DESeq2 (v1.44.0, Love et al., 2014). It was performed for each plant-fungal interaction comparing inoculated vs non-inoculated plants at early and late developmental stages, as explained in the experimental design (**Table S1**). Condition-specific differentially expressed genes (DEGs) with a false discovery rate (FDR) < 0.01 and a log2 fold-change (Log2FC) ≥ |1|, were identified for each pairwise condition. Volcano plots were used to illustrate and highlight DEGs. To consider a putative gene as small secreted protein (SSPs), the following criteria were used: 1) genes encoding a protein less than 300 aa; 2) genes with a predicted signal peptide (> 0.5); 3) orphan genes without known domains and homology with other species.

## Data statement

All the article’s supporting data and materials can be accessed in the main body and in supplementary files. RNAseq raw datasets of poplar interaction can be accessed from Short Read Archive (SRA) database (https://www.ncbi.nlm.nih.gov/sra), with the accession numbers indicated in **Table S1**.

## Acknowledgements

This research was sponsored by the Plant Microbe Interfaces Scientific Focus Area (http://pmi.ornl.gov) in the Genomic Science Program of the Office of Biological and Environmental Research in the U.S. Department of Energy Office of Science (JEMG, FM, CVF, AK). The Oak Ridge National Laboratory is managed by UT-Battelle in the U.S.DOE Office of Science (DE-AC05-589 00OR22725). JEMG (123/MTAI/22) is grateful to the funding institutions for their contracts with the University of Murcia financed by the Ministry of Universities through the NextGenerationEU funds of the European Union and the Recovery, Transformation and Resilience Plan of the Spanish Government through the Programme for the Requalification of the Spanish University System during the three-year period 2021-2023. MFP’s doctoral fellowship and her research visit at INRAE Grand Est-Nancy were supported by the Brazilian agencies: Fundação de Amparo à Pesquisa do Estado de Minas Gerais (FAPEMIG), Coordenação de Aperfeiçoamento de Pessoal de Nível Superior (CAPES), and Conselho Nacional de Desenvolvimento Científico e Tecnológico (CNPq). Part of the research visit of MFP and the BLACKSECRET project to AK and MP were financed by the Laboratory of Excellence ARBRE (ANR-11-LABX-0002-01) and by WSL. JR was an Agreenskills Marie Sklodowska Curie fellow co-funded by the European Commission (CoFUND-FP7-267196)). The sequencing project (proposal 10.46936/10.25585/60001022) was conducted by the U.S. Department of Energy Joint Genome Institute (https://ror.org/04xm1d337), a DOE Office of Science User Facility, and is supported by the Office of Science of the U.S. Department of Energy under Contract No. DE-AC02-05CH11231.

## Short legends for Supporting Information

**Table S1.** RNAseq datasets used in this study.

**Table S2.** Differentially expressed poplar genes (Log2 fold change > |1| and False Discovery Rate < 0.01) in ECM roots vs non-inoculated roots at early stage. (A) *L. bicolor*; (B) *C. geophilum*; (C) *P. microcarpus*.

**Table S3.** Differentially expressed poplar genes (Log2 fold change > |1| and False Discovery Rate < 0.01) in ECM roots vs non-inoculated roots at mature stage. (A) *L. bicolor* (1); (B) *L. bicolor* (2); (C) *C. geophilum*; (D) *A. muscaria*; (E) *P. microcarpus*.

**Table S4.** ECM core gene regulon. (A) 125 poplar gene regulon in response to ECM symbiosis at early stage. (B) 138 poplar gene regulon in response to ECM symbiosis at mature stage.

**Table S5.** Gene Ontology enrichment in the 125 ECM core gene regulon at early stage. (A) GO terms significantly enriched (p < 0.01). (B) Differentially expressed genes belonging to significantly enriched families.

**Table S6.** Gene Ontology enrichment in the 138 ECM core gene regulon at mature stage.

(A) GO terms significantly enriched (p < 0.01). (B) Differentially expressed genes belonging to significantly enriched families.

**Table S7.** Differentially expressed poplar genes (Log2 fold change > |1| and False Discovery Rate < 0.01) in inoculated roots vs non-inoculated roots at early stage. (A) *P. tinctorius*; (B) *S. citrinum*.

**Table S8.** Differentially expressed poplar genes (Log2 fold change > |1| and False Discovery Rate < 0.01) in inoculated roots vs non-inoculated roots at mature stage. (A) *P. tinctorius*; (B) *S. citrinum*.

**Table S9.** DEG count of Pathogenesis-related genes and Serine proteinase inhibitors of poplar roots in co-culture with *L. bicolor*, *C. geophilum*, *P. microcarpus*, *P. tinctorius* and *S. citrinum* at different stages.

**Table S10.** Differentially expressed fungal genes (Log2 fold change > |1| and False Discovery Rate < 0.01) in co-culture with *P. tremula x alba* vs free living mycellium at early stage. (A) *P. microcarpus*; (B) *P. tinctorius*; (C) *S. citrinum*.

**Table S11.** Differentially expressed fungal genes (Log2 fold change > |1| and False Discovery Rate < 0.01) in co-culture with *P. tremula x alba* vs free living mycelium at mature stage. (A) *P. microcarpus*; (B) *P. tinctorius*; (C) *S. citrinum*.

**Figure S1.** Volcano plots representing differentially expressed genes of ECM compatible interactions at early and mature stages of plant-fungus interaction.

**Figure S2.** Sequence alignment of poplar germin-like genes containing cupin domain and belonging to the ECM core gene regulon.

**Figure S3.** Sequence alignment of poplar Pollen Ole e1 genes containing proline rich extensin signatures and belonging to the ECM core gene regulon.

**Figure S4.** Volcano plots representing differentially expressed genes of non-compatible interactions at early and mature stages of plant-fungus interaction.

**Figure S5.** Transcripts per Million reads (TPM) for genes of the common symbiosis pathway and related to AM symbiosis.

**Figure S6.** Sequence alignment of *Sclerodermataceae* small secreted proteins homologs to *Pm683008*.

**Figure S7.** Quality control of poplar RNAseq analyses of ECM compatible interactions at both early and mature stages.

**Figure S8.** Quality control of poplar RNAseq analyses of non-compatible interactions at both early and mature stages.

**Figure S9.** Quality control of *P. microcarpus* RNAseq analyses at both early and mature stages.

**Figure S10.** Quality control of *P. tinctorius* RNAseq analyses at both early and mature stages.

**Figure S11.** Quality control of *S. citrinum* RNAseq analyses at both early and mature stages.

