## Supplementary figures and images for "Comparative transcriptomics uncovers plant and fungal genetic determinants of mycorrhizal compatibility"

### Supplemental Figure 1

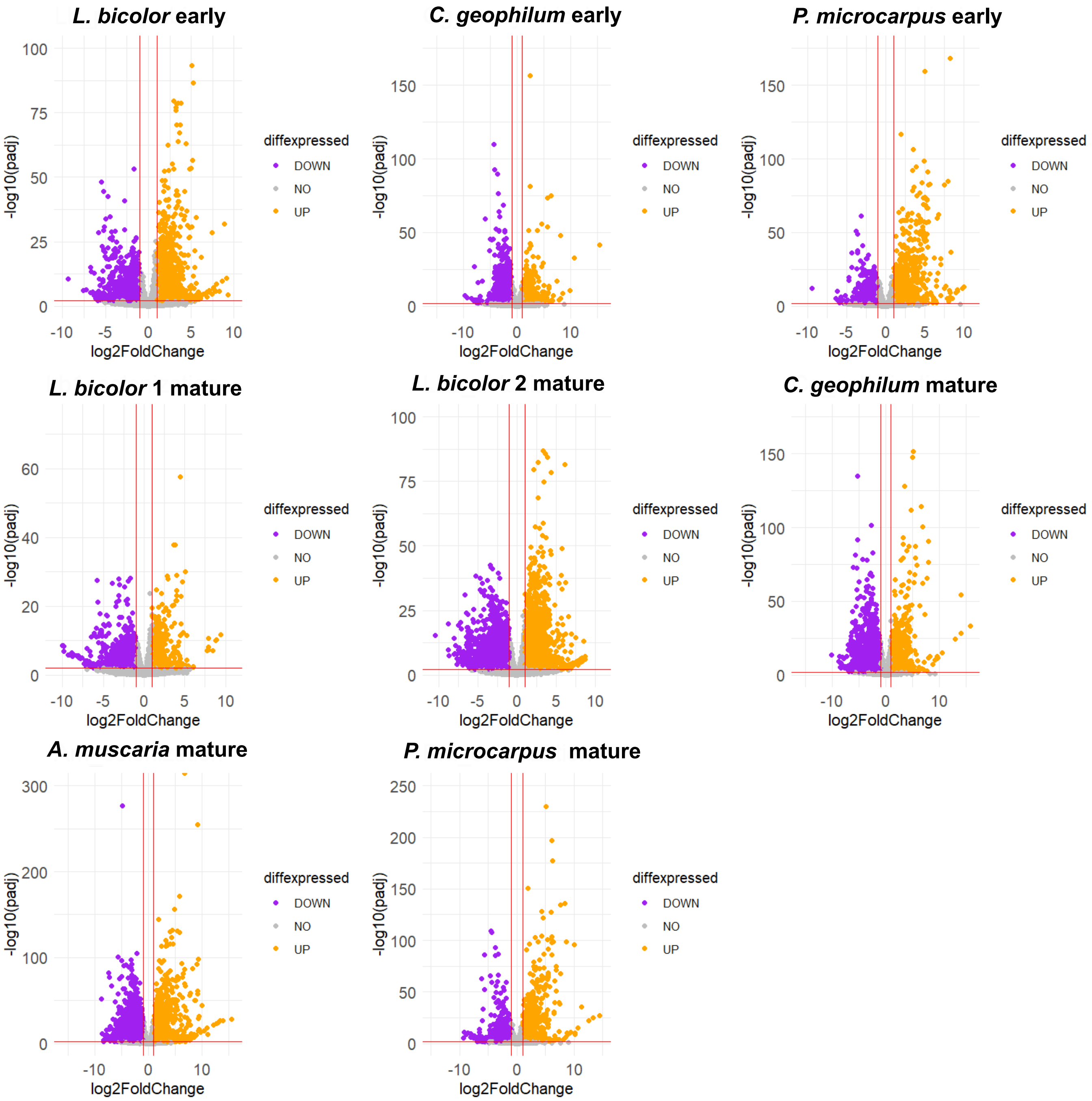

### Supplemental Figure 2

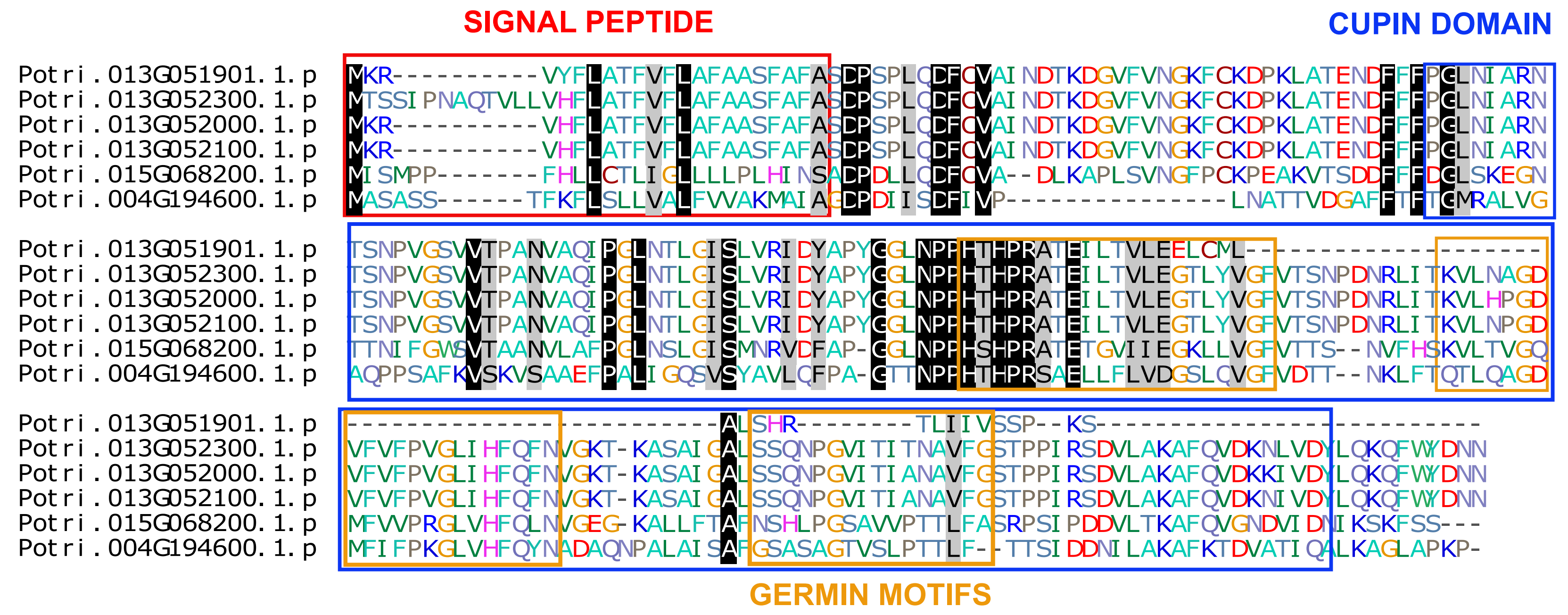

### Supplemental Figure 3

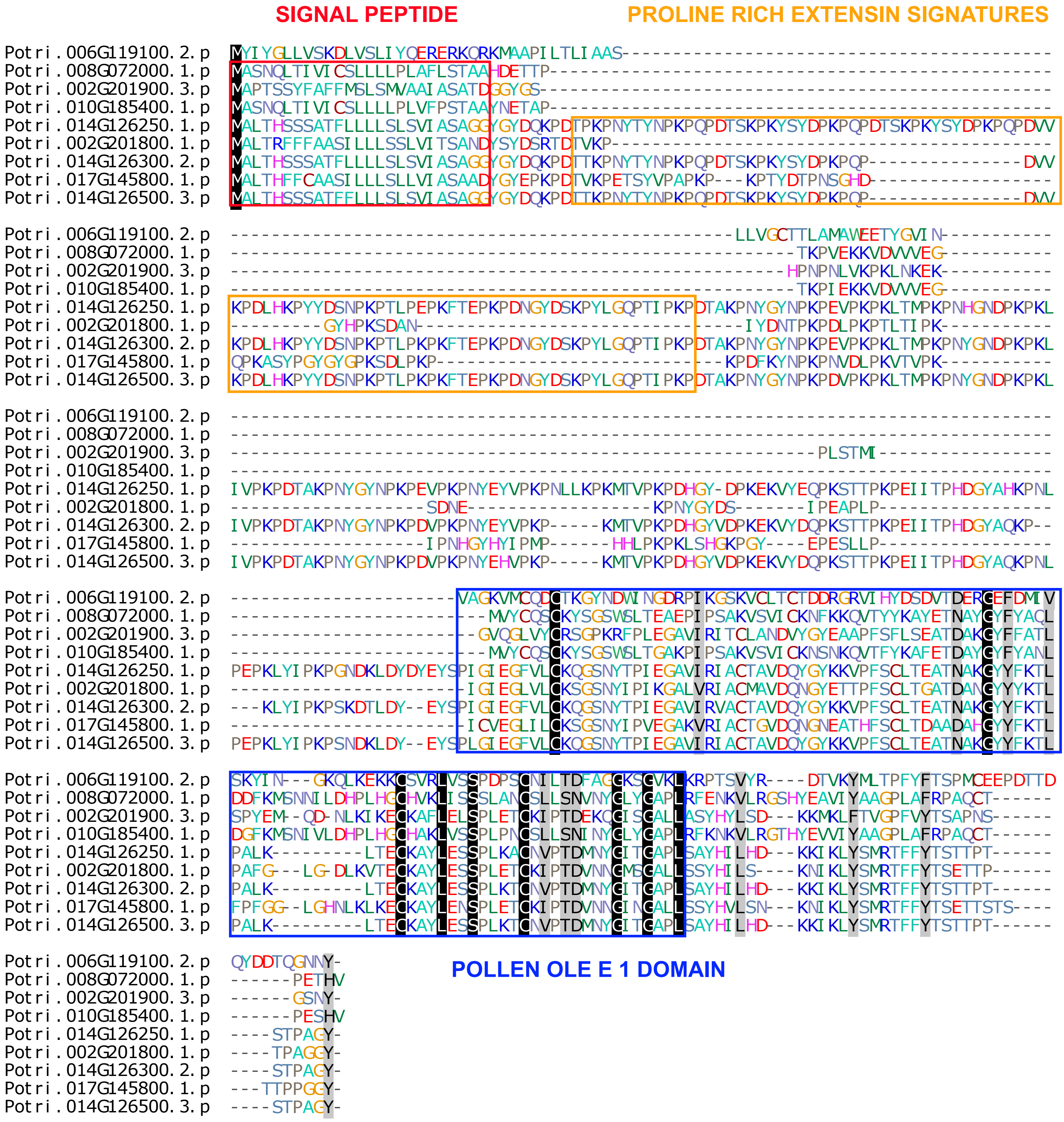

### Supplemental Figure 4

***S. citrinum* mature**

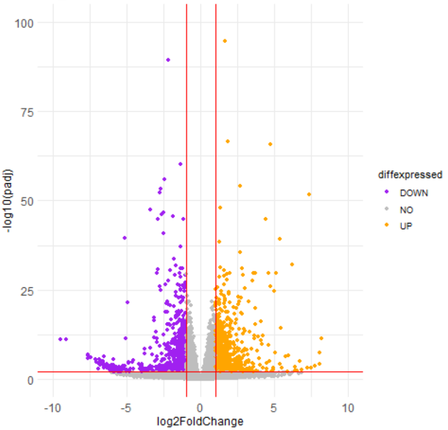


***P. tinctorius* early**

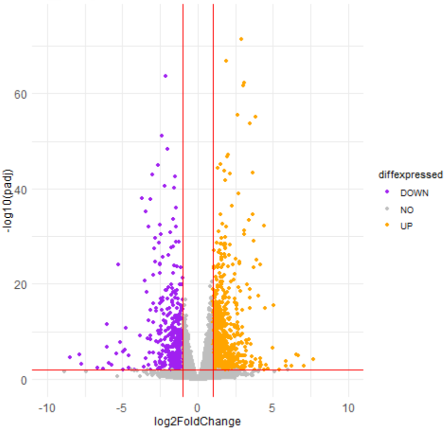


***S. citrinum* early**

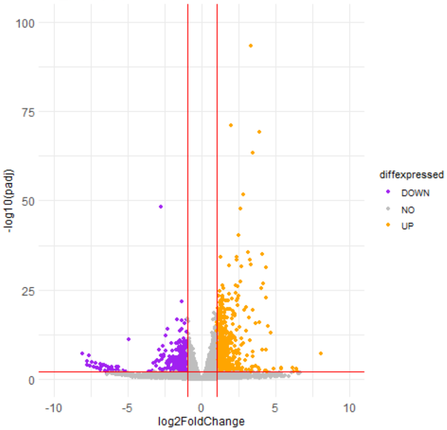


***P. tinctorius* mature**

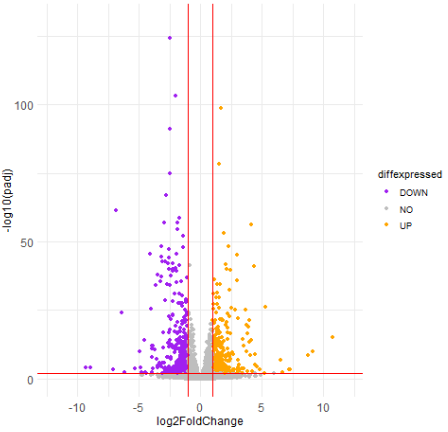

### Supplemental Figure 5

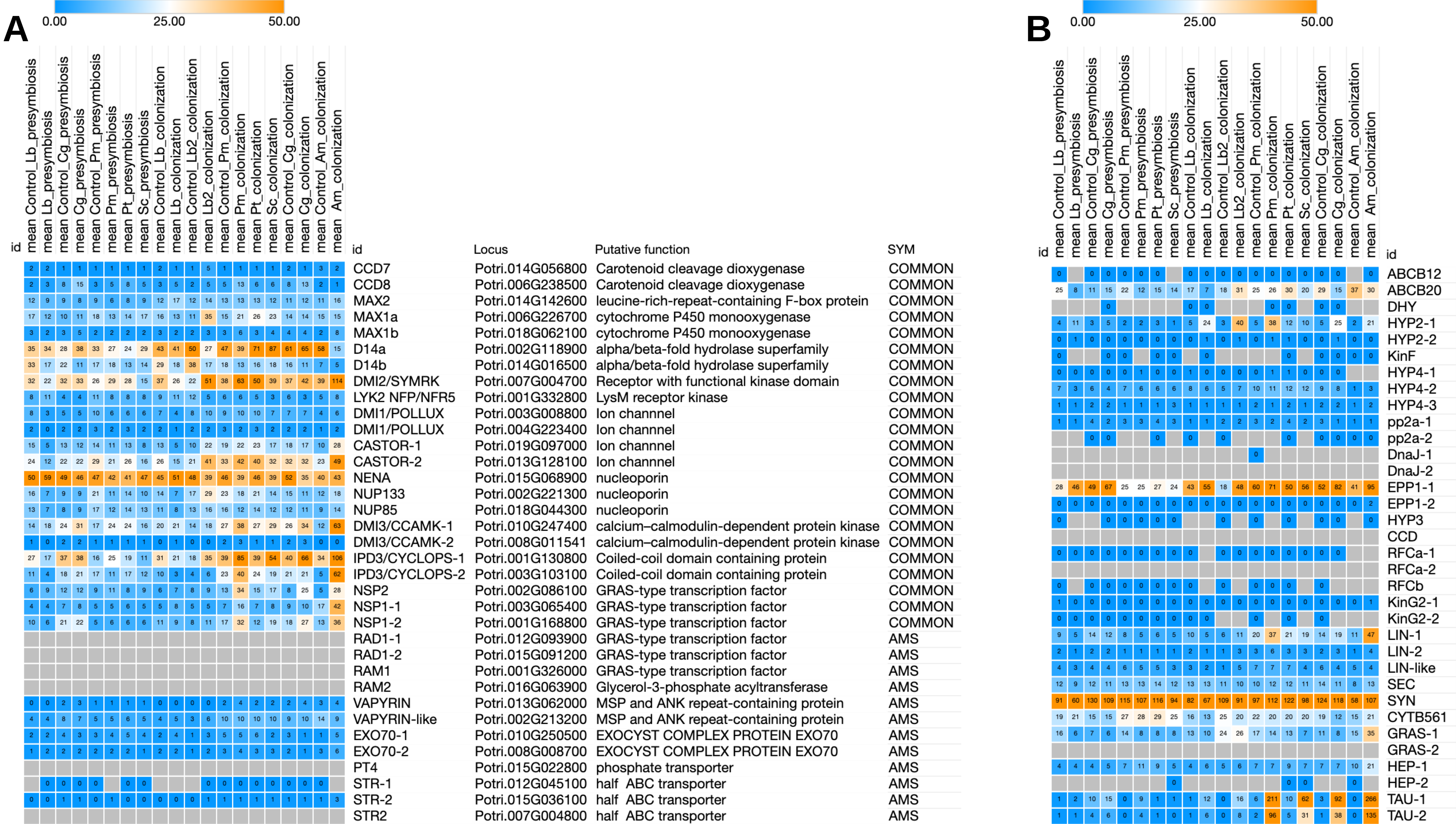

### Supplemental Figure 6

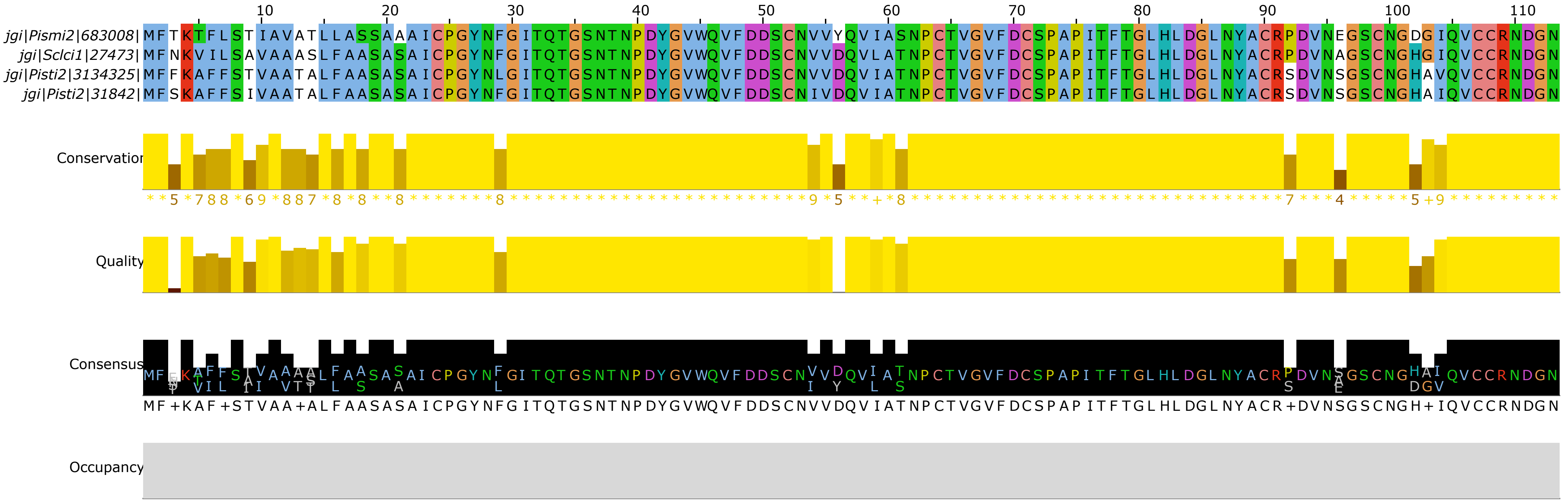

### Supplemental Figure 7

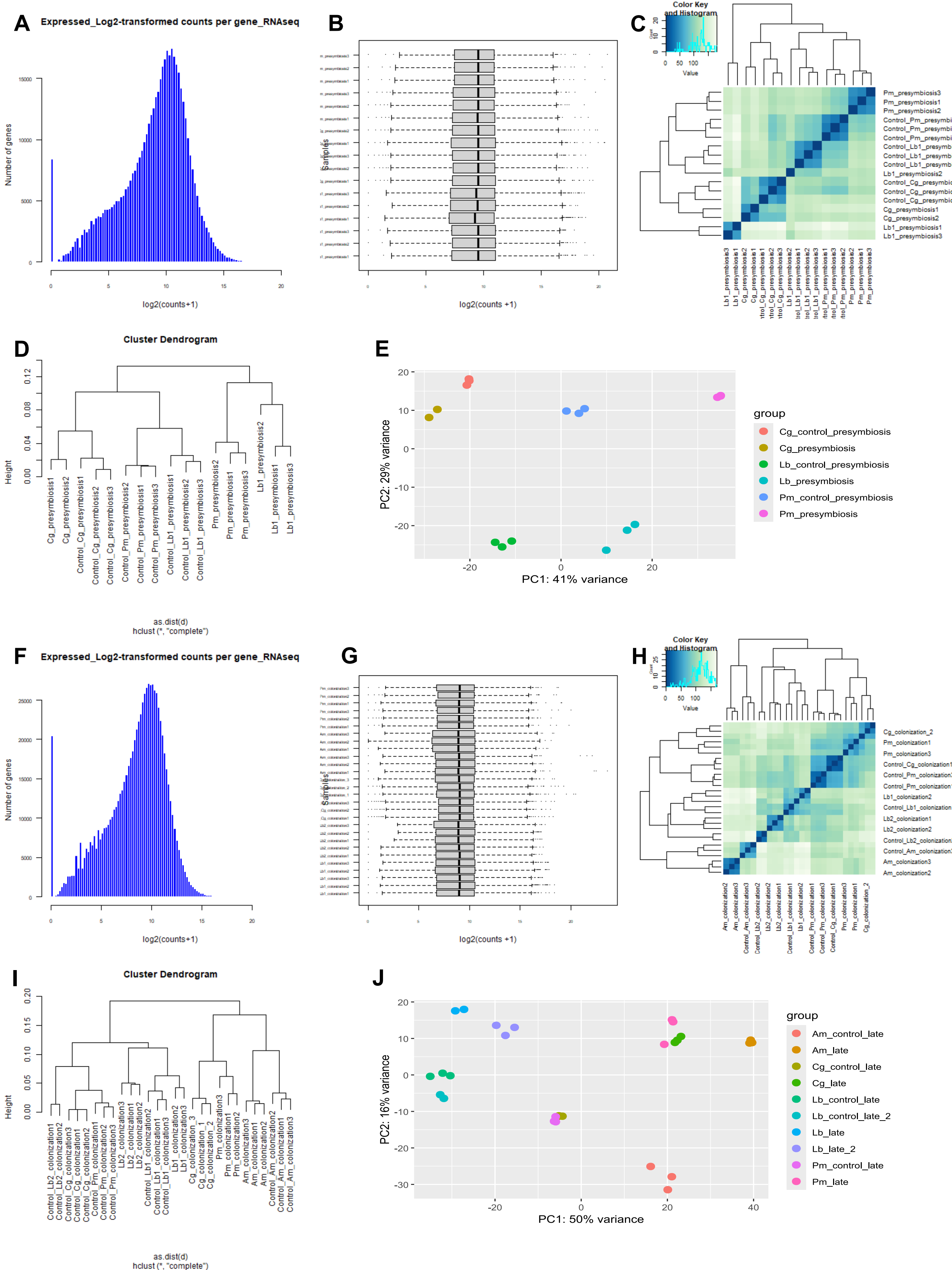

### Supplemental Figure 8

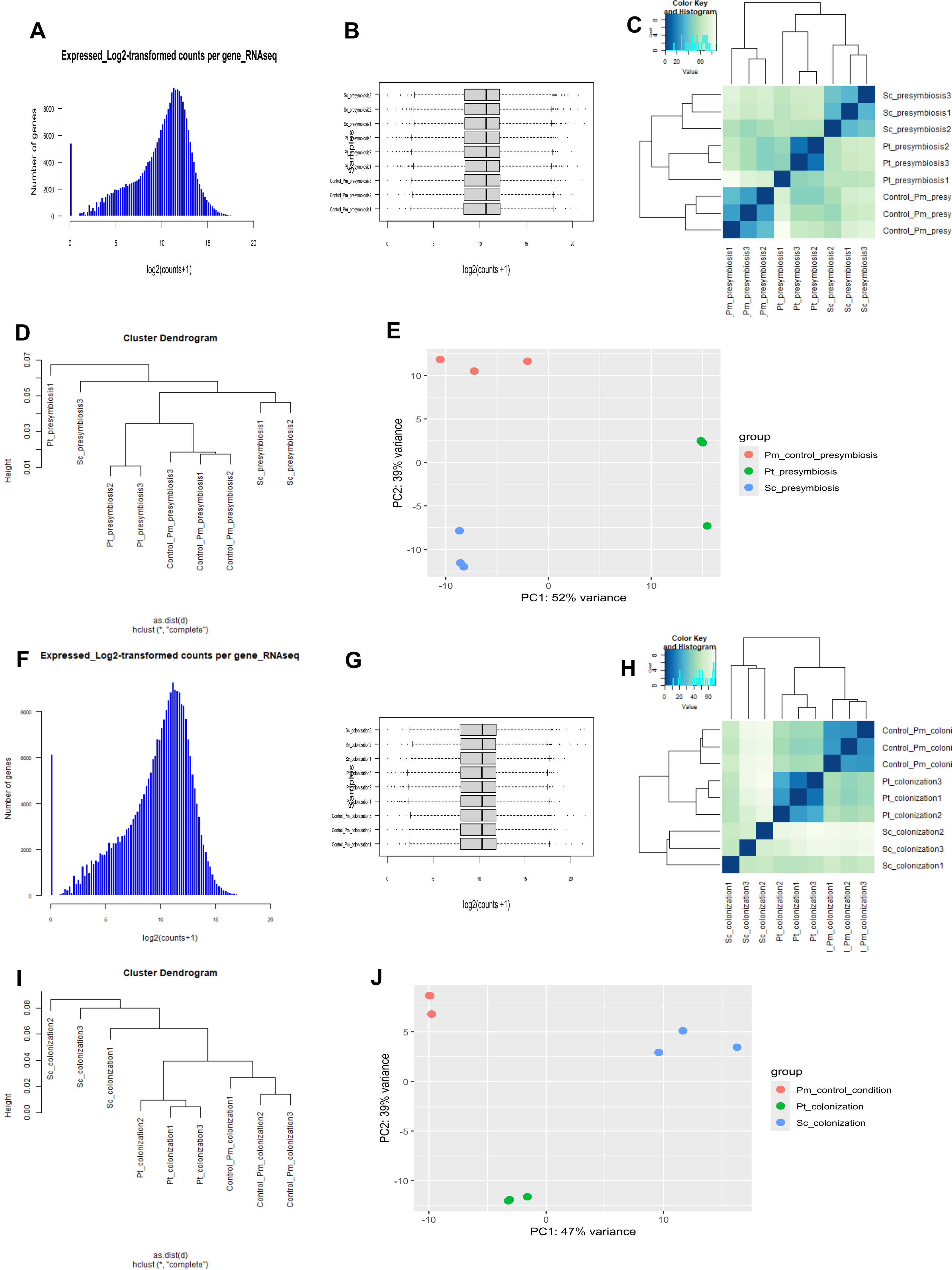

### Supplemental Figure 9

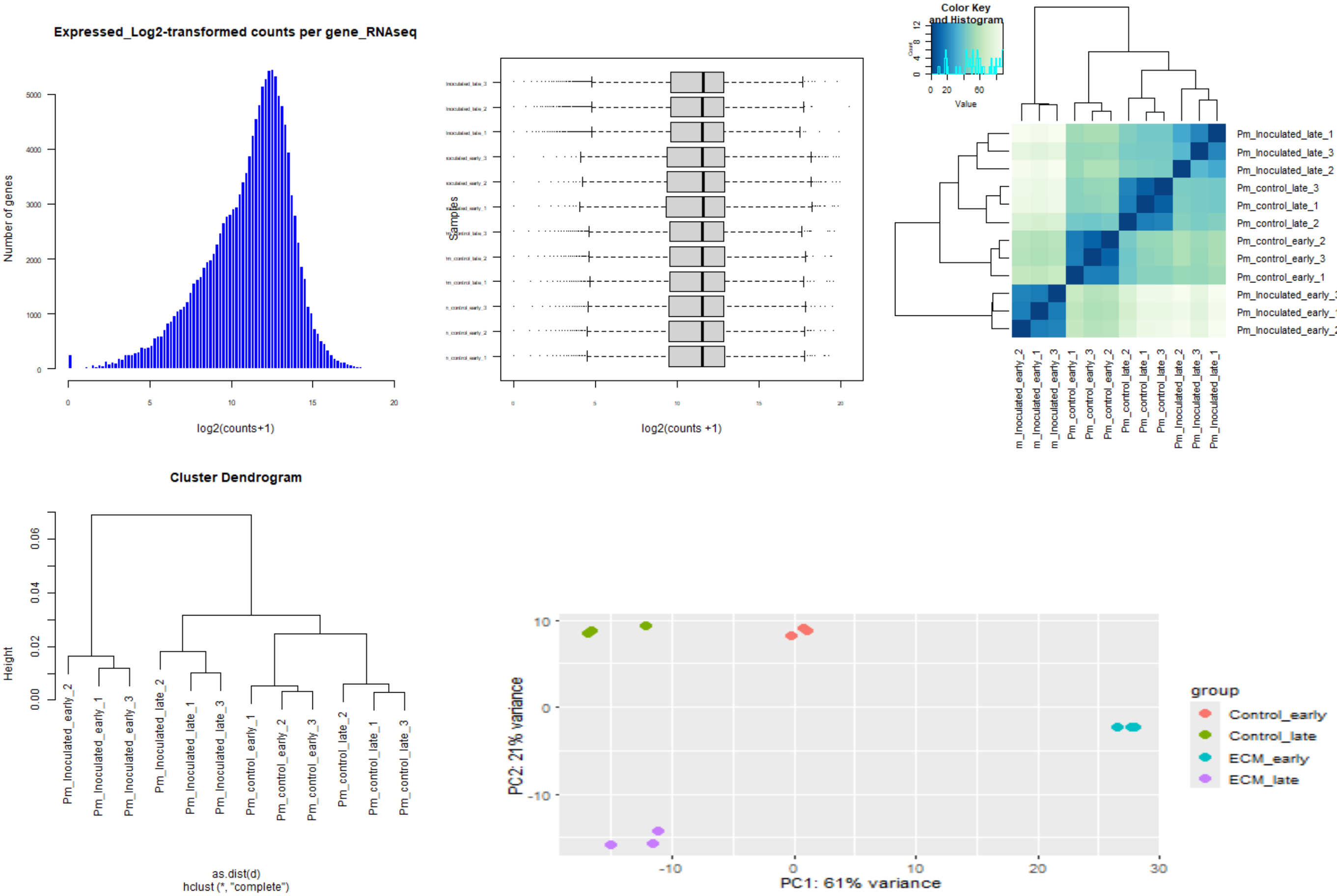

### Supplemental Figure 10

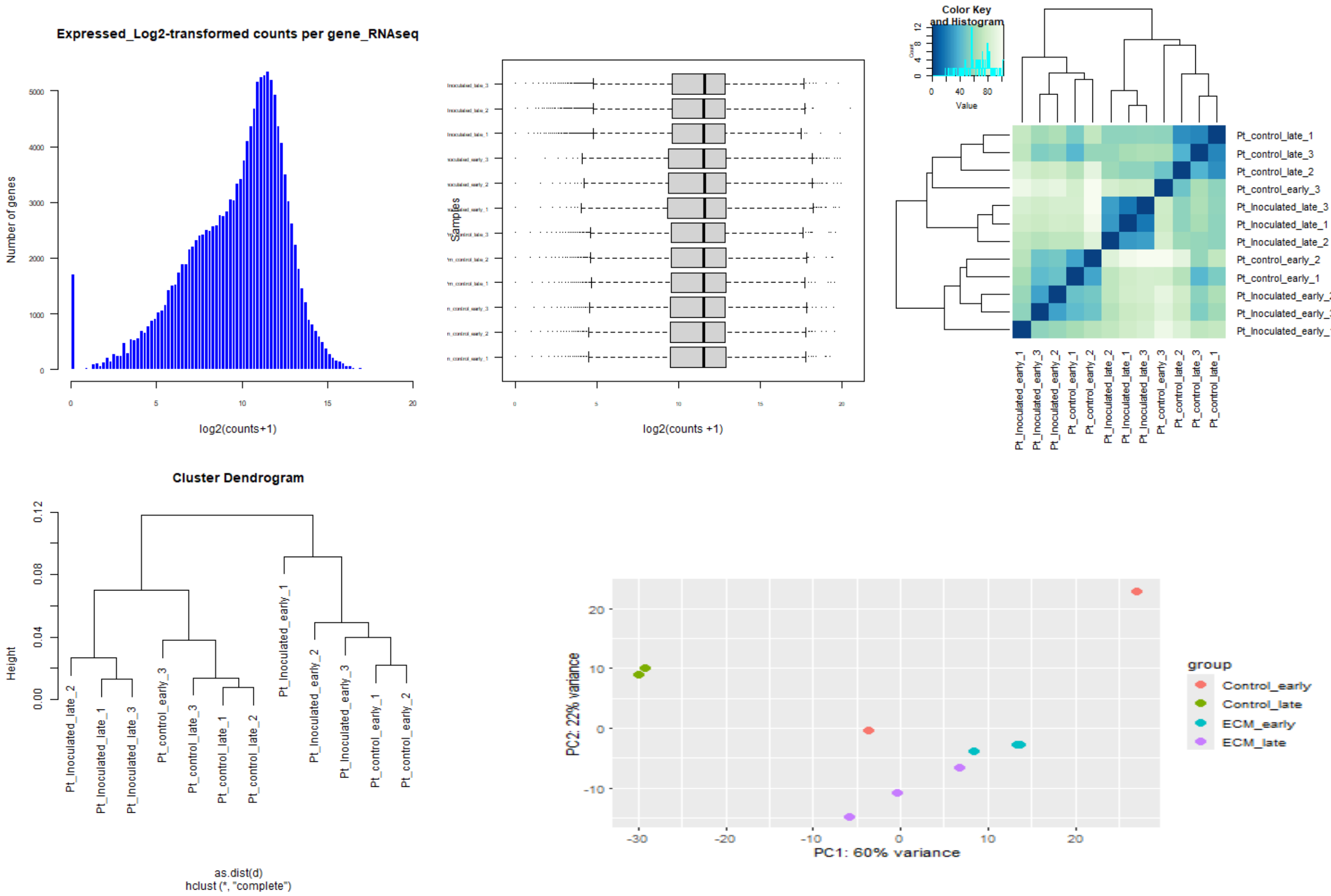

### Supplemental Figure 11

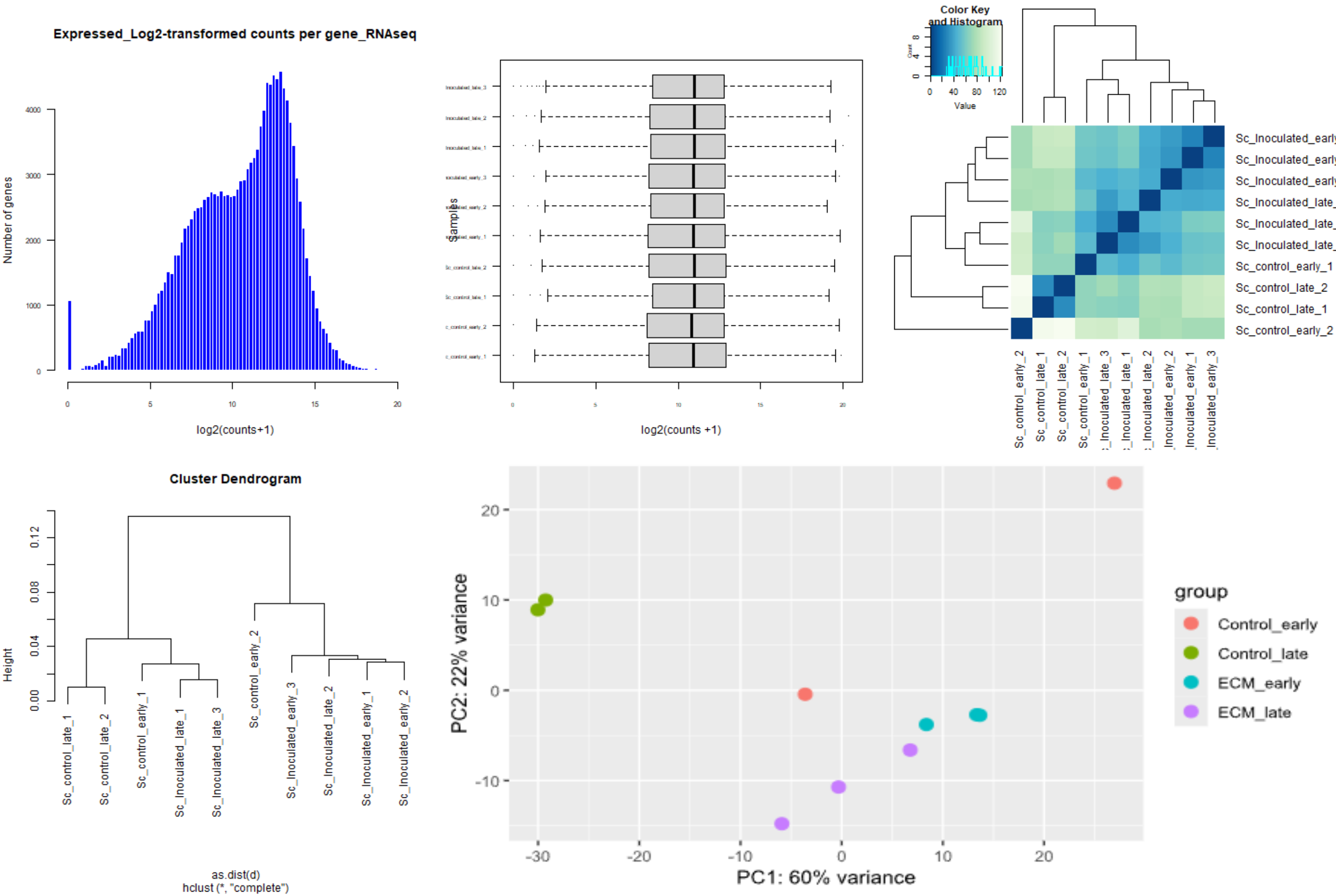
